# Selection for early reproduction leads to accelerated aging and extensive metabolic remodeling in *Drosophila melanogaster*

**DOI:** 10.1101/2024.06.28.601037

**Authors:** David L. Hubert, Kenneth R. Arnold, Zachary S. Greenspan, Anastasia Pupo, Ryan D. Robinson, Valeria V. Chavarin, Thomas B. Barter, Danijel Djukovic, Daniel Raftery, Zer Vue, Antentor Hinton, Melanie R. McReynolds, Benjamin R. Harrison, Mark A. Phillips

## Abstract

Experimental evolution studies that feature selection on life-history characters are a proven approach for studying the evolution of aging and variation in rates of senescence. Recently, the incorporation of genomic and transcriptomic approaches into this framework has led to the identification of hundreds of genes associated with different aging patterns. However, our understanding of the specific molecular mechanisms underlying these aging patterns remains limited. Here, we incorporated extensive metabolomic profiling into this framework to generate mechanistic insights into aging patterns in *Drosophila melanogaster*. Specifically, we characterized metabolomic change over adult lifespan in populations of *D. melanogaster* where selection for early reproduction has led to an accelerated aging phenotype relative to their controls. Using these data we: i) evaluated evolutionary repeatability across the metabolome; ii) assessed the value of the metabolome as a predictor of “biological age” in this system; and iii) identified specific metabolites associated with accelerated aging. Generally, our findings suggest that selection for early reproduction resulted in highly repeatable alterations to the metabolome and the metabolome itself is a reliable predictor of “biological age”. Specifically, we find clusters of metabolites that are associated with the different rates of senescence observed between our accelerated aging population and their controls, adding new insights into the metabolites that may be driving the accelerated aging phenotype.

**Significance:** While experimental evolution studies featuring *Drosophila melanogaster* have generated significant insights into the forces that shape aging and life history patterns, more recent efforts incorporating genomic and transcriptomic data have had comparatively little success identifying the molecular mechanisms underlying these patterns. Here we work to incorporate molecular phenotyping into this general framework as a way forward. Specifically, we characterize how the metabolome changes with age in populations of *D. melanogaster* where hundreds of generations of selection for early reproduction have led to enrichment for an accelerated aging phenotype. By comparing to control populations, we show that the metabolome does appear to capture true signals of “biological age” and provides a new avenue for understanding the factors that underlie complex trait variation in real populations.

## Introduction

Experimental evolution studies using *Drosophila melanogaster* populations have been a continual source of insight into the factors that underlie differences in longevity and rates of senescence between individuals. As predicted by the evolutionary theory of aging, selection for early reproduction reliably produces populations enriched for individuals that develop quickly and die young, whereas selection for postponed reproduction has the opposite effect (Rose 1984; Luckinbill et al. 1984; Chippindale 1997; Rose et al. 2010). Studying these populations has in turn generated many novel insights into the physiological trade-offs associated with different rates of senescence. For instance, while populations selected for early reproduction develop quickly and have high early life fecundity, they also exhibit reductions in egg viability (Chippindale et al. 1997), stress resistance (Kezos et al. 2023), lifetime fecundity, and lifespan (Burke et al. 2016). The goal of the current project is to build on this work to better understand the molecular underpinnings of this “accelerated aging” phenotype.

Over the past decade, the combination of experimental evolution and next-generation sequencing technologies has emerged as a powerful tool for studying the genetics of complex traits. Major findings from studies that analyze quantitative traits, such as longevity, stress resistance, and body size, have consistently shown that the underlying genetic architecture is complex, often involving hundreds of genes (e.g. Burke et al. 2010; Turner et al. 2011; Fabian et al. 2018; Tobler et al. 2015; Barter et al. 2019; Kezos et al. 2019). Studies have also found that many of these traits may involve genes with significant pleiotropic interactions (Greenspan et al. 2024), and suggest genetic redundancy is a common feature of complex trait genetic architecture (Hardy et al. 2018; Barghi et al. 2019). Because of this inherent complexity, direct insights into the molecular mechanisms underlying trait variation have been modest. With respect to aging and longevity, functional conclusions are largely limited to enrichment for gene ontology terms broadly related to neurological development, immune function, and metabolism (Carnes et al. 2015; Fabian et al. 2018, Barter et al. 2019). In an effort to move beyond the finding that complex traits are highly polygenic, some have turned to metabolomics as a path forward (e.g. Phillips et al. 2022; Cavigliasso et al. 2023).

Metabolomics involves the profiling of metabolites, molecules <2000 Da, collectively referred to as the metabolome. Because metabolites constitute the functional and energetic building blocks of all life, metabolomics provides a potential link between genotype and phenotype (Harrison et al. 2020). Among the -omic layers of systems biology (e.g. transcriptome, proteome, etc.) the metabolome is thought to closely reflect organismal phenotypes (Dettmer et al. 2007; Hoffman et al. 2014). While genomic, transcriptomic, and proteomic signals can be complicated by epigenetic and posttranslational modifications, it is thought that the downstream positionality of the metabolome provides a marker more representative of the observed phenotype (Patti et al. 2012). The metabolome of flies is highly dynamic with age and is sensitive to effects on longevity from environmental conditions, genetic variation, and pharmacological interventions (Avanesov et al. 2014; Hoffman et al. 2014; Laye et al. 2015; Jin et al. 2020; Phillips et al. 2022; Wang et al. 2022). In light of this, recent efforts have been made to include metabolomic phenotyping into the evolutionary biology framework (Harrison et al. 2022; Phillips et al. 2022; Erkosar et al. 2024).

With respect to characterizing age sensitive effects, Zhao et al. (2022) recently used metabolomic data to build a multivariate age-prediction model colloquially referred to as a “metabolomic clock”. As with other -omic clocks such as epigenetic clocks, the age prediction is not fully accurate (Rutledge et al. 2022; Parrott and Bertucci 2019). Indeed, the deviation of the age prediction from the fly metabolome clock from chronological age of the sampled flies, the flies so-called ‘age acceleration’, correlated with mortality rate and the remaining life expectancy of isogenic flies (Zhao et al., 2022). That is, fly strains whose predicted age was younger than their chronological age tended to live longer than flies whose predicted age was greater than their chronological age. Thus, the metabolome clock connects patterns of age in the metabolome to the rise in mortality risk that defines biological aging. Using clock models could provide insight into the molecules associated with the relative pace of chronological and biological age, where the former is simply a measure of the passing of time, and the latter is a relative measure of senescence (Jylhävä et al. 2017). In an experimental evolution context, such clock models could allow us to test hypotheses about the aging patterns that result from selection on life history (Parrott and Bertucci 2019).

Here, we applied metabolomic analysis to a set of ten replicate fruit fly populations where hundreds of generations of selection for early reproduction (Figure 1) has led to enrichment for an accelerated aging phenotype where flies develop quickly and die young relative to controls. This accelerated aging phenotype presents as a disconnect between chronological and biological age when compared to the ancestral population. Previous work in this system has consistently demonstrated the evolutionary repeatability of this accelerated aging signal with high levels of genomic (Graves et al. 2017), transcriptomic, (Barter et al. 2019), and phenotypic convergence (Burke et al. 2016; Kezos et al. 2023) between newly derived and long-standing populations subjected to the same selection regime. This rapid convergence has been attributed to the hypothesis that adaptation in these populations is primarily fueled by shifts in common genetic variants present in the ancestral population that this system was derived from.

**Figure 1.**
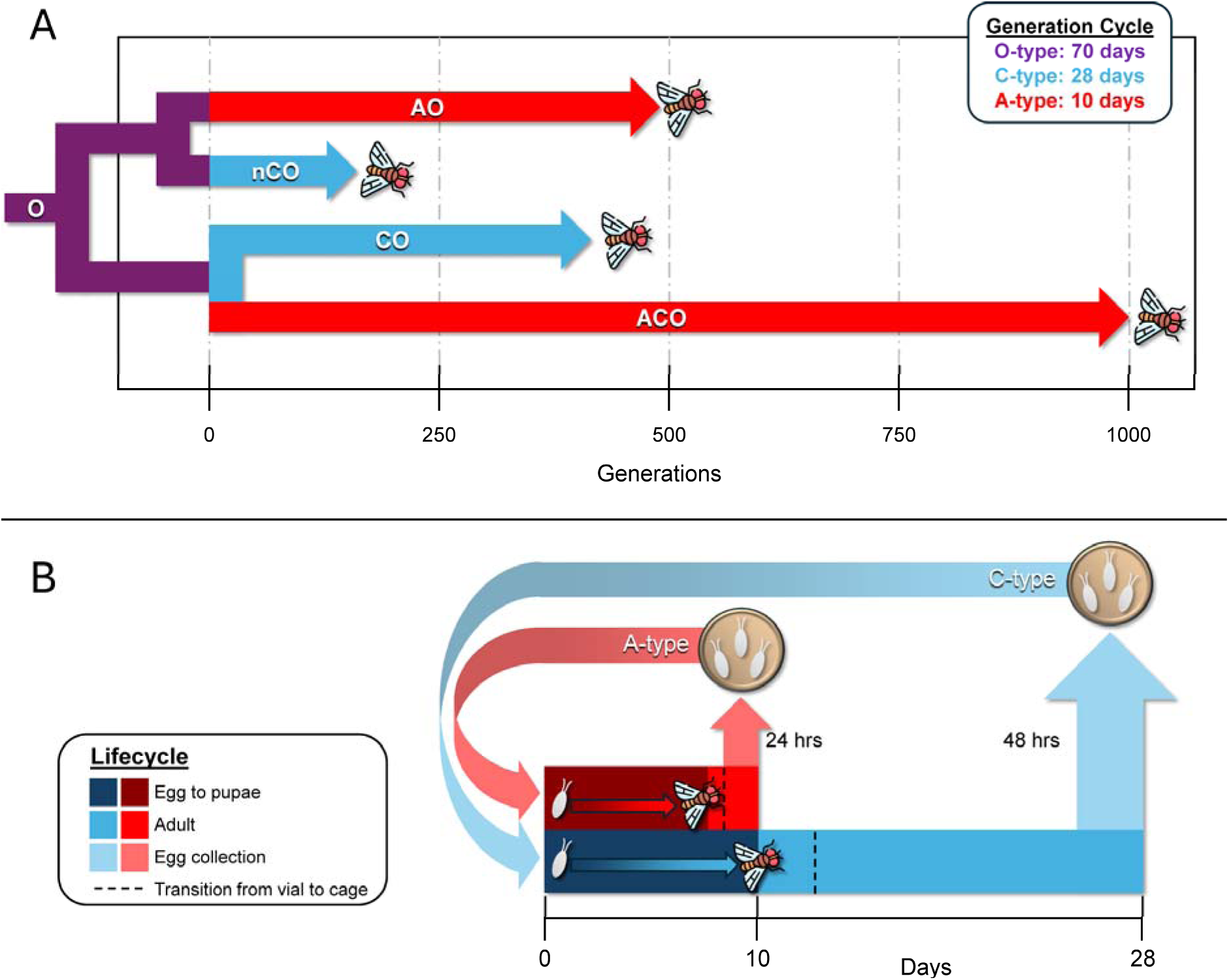
Recent phylogeny and selection regimes for *D. melanogaster* populations of interest. A: Abbreviated diagram of the evolutionary relationship between the populations of interest. The ancestral population for all populations was maintained on a 70-day generation cycle (O-types), both the long-standing (CO) and recently derived (nCO) C-type populations have been maintained on 28-day generation cycles, and both the long-standing (ACO) and recently derived (AO) A-type populations have been maintained on 10-day generation cycles. B: Experimentally controlled reproductive windows used to maintain the two life-history regimes: Accelerated (A), with a 10-day reproductive cycle; and Delayed (C), with a 28-day reproductive cycle. Short reproductive windows, either 24 hours for Accelerated or 48 hours for Delayed, were used to enforce reproductive cycle length.

In Phillips et al. (2022), we compared metabolomic differentiation patterns between samples of 21-day-old flies (age from egg) from the accelerated aging (A-type) populations and their controls (C-type) and found evidence of altered tricarboxylic acid (TCA) cycle activity, carbohydrate metabolism, and neurological function in the A-type populations. Because age-specific mortality rates are known to greatly diverge between the fly populations by day 21, we concluded that the observed differences in metabolic activity were a potential driver of the aging and longevity differences observed between the selection regimes. One possibility is that these differences were due to metabolic remodeling, similar to the differences between reproductive and non-reproductive flies (Koliada et al. 2020; Rodrigues et al. 2023). Such remodeling in the A-type populations could meet energetic and nutritional demands associated with fast development and early reproduction and come at the expense of somatic maintenance. However, our ability to make a definitive statement was limited as Phillips et al. (2022) included only a single age class and did not establish if this metabolomic difference reflected a difference in biological aging. In the present study, we seek to remedy this shortcoming and derive even deeper mechanistic insights by testing for accelerated biological age and characterizing how metabolomic profiles change over time in the A and C-type regimes by sampling at multiple age classes (Figure 2).

**Figure 2.**
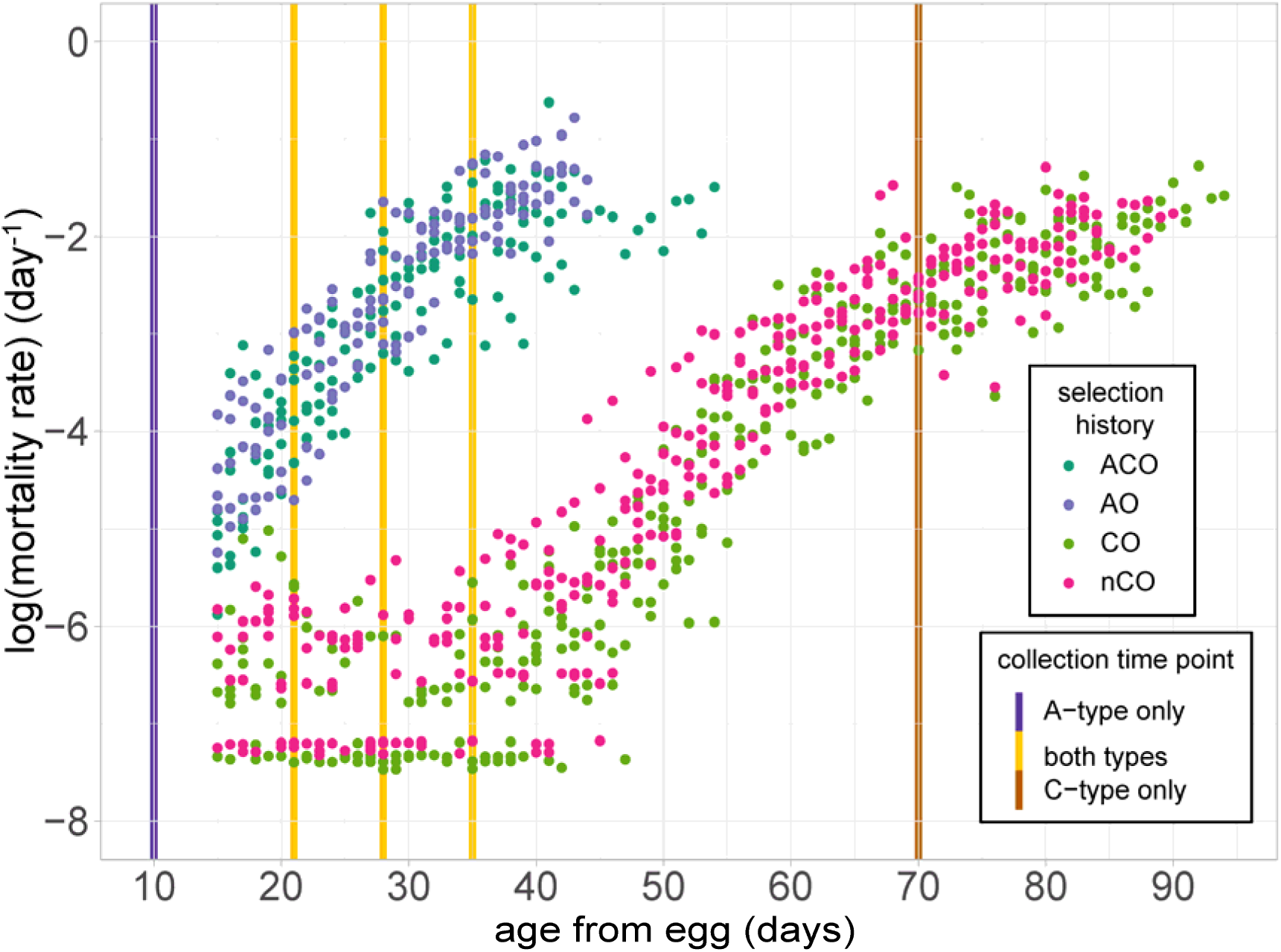
Age-specific mortality and metabolomic sampling design. Female mortality rates for 20 cohorts of A-type (blue and purple) and C-type (pink and green) populations (See methods for calculation). Selection history is further indicated by color, with ACO and CO representing long-standing populations and AO and nCO representing recently derived populations. Each vertical bar represents the collection timepoint in either A-type (blue), C-type (orange) populations, or both (yellow).

We hypothesize that the metabolome is connected to the physiological tradeoffs that accompany adaptation to the A-type selection regime. If true, we predicted that i) metabolomic profiles should rapidly converge for populations subjected to the same selective pressures, ii) a metabolomic clock should not only capture signals of chronological age, but also reflect differences in biological age associated with the response to different selective pressures, and iii) characterization of metabolite abundance over time should reveal specific metabolites associated with the accelerated aging phenotype in flies under the A-type selection regime, potentially providing mechanistic insights.

To test these predictions and further explore the data, we had three primary objectives:

Objective 1) To test our first predictions, we compared the metabolomes of long-standing and recently derived populations subjected to the same selection regime to evaluate levels of metabolomic convergence between them. If the known life history trade-offs that characterize the response to this selection in this system have a clear signal in the metabolome, newly derived and long-standing populations should show a high degree of metabolomic convergence within each respective regime.

Objective 2) To test our second prediction, we created separate metabolomic clocks using data from A or C-type regimes. As with previous studies, we expect that these models will capture signals associated with age. In other studies, the deviation of the clock’s prediction from the chronological age of the sampled flies reflected mortality risk within the studied population. In this setting however, we use such age acceleration to test our hypothesis about the relative biological ages of flies from these two regimes. We expect that, relative to the metabolome of the C-type populations, the A-type population metabolome will appear ‘older’. Similarly, we predict that a clock model based on the metabolome of the A-type populations will predict younger biological age for the metabolome of the C-type populations.

Objective 3) To test our third prediction, we assessed the similarity of the metabolome of all samples by hierarchical clustering. We expect that an accelerated aging phenotype will be represented in the metabolome via the presence of clusters of metabolites whose abundance shows greater similarity between the old C-type and the younger A-type populations. We then use these clustered metabolites to generate insights into the metabolic mechanisms underlying the accelerated aging phenotype observed in flies from the A-type regime.

## Results

### Metabolomic profiling

Targeted metabolomic profiling included a panel of 361 target metabolites, 210 of which were detected during LC-MS profiling, with 202 including values for all samples (see Table S1 for full details). Metabolite abundance values were normalized (Methods) for further analyses (Table S2).

A preliminary principal components analysis was conducted to look for general clustering by selection regime, age, and selection history (Figure S1). The common ages sampled for both the A-type and C-type populations (21, 28, and 35 days from egg) showed separation along both the PC1 and PC2 axes. The 9-day-old samples from A-type populations clustered separately from all other samples along both PC axes, showing a greater degree of differentiation for these samples than the rest. This striking difference in metabolomic profile of the 9-day-old samples is likely due to the residual effects of recent eclosion (<24 hours), which can have large, short-term impacts on energy reserves (Erkosar et al. 2024). The 70-day-old samples from C-type populations were the only samples that clustered more closely with the opposing selection type, showing overlap with samples from the A-type populations at ages 28- and 35-days-old. Samples from the common ages (21, 28, and 35 days) clustered within selection regime, with some separation by age within their respective selection regimes. These patterns were consistent when selection history (recently derived or long-standing) was factored (Figure S1).

#### Objective 1: Metabolomic Convergence Within Selection Regime

Because of the intimate connection between metabolome and physiology, we hypothesize that the among-regime convergence in mortality among the more-recently derived and the long-standing selection lines is paralleled by convergence in their metabolome. We tested convergence among the metabolome of more-recently derived and the long-standing populations in two ways, both of which test for *divergence* of the metabolome. In this way, we are asking if the metabolome of recent and long-standing populations subject to the same selection regime differ significantly. Such effects would be contrary to our hypothesis. First, by testing for effects of selection history (recent v. long-standing) within each regime on the multivariate metabolome, and second, by testing the accuracy of predictive models trained on the metabolome of long-standing populations to predict the regime of more recently derived populations based on their metabolome.

We used principal components analysis to reduce the dimensionality of the metabolome data from on the 21-, 28- and 35-day from egg timepoints on which samples from both regimes A and C were sampled (Table S3). Among the first four PCs, which together explain 60.6% of the variance, PCs 1 and 3 showed strong regime-dependent effects of age, while PCs 2 and 4 were patterned by age, and PC3 by main effects of regime (ANOVA P<5% after Bonferroni correction, Methods, Figure 3). We tested for divergence on the multivariate metabolome as interaction effects of history within regimes. We found no evidence of regime x history interaction on any of the PCs, either as main effects on the PC (PC ∼ regime x history), or on the trajectory of PCs over age (PC ∼ regime x history x age; P>5%, Figure 3, Methods).

We then tested the accuracy of a supervised multivariate predictive model trained on only the metabolome data of the long-standing populations collected on days 21, 28 and 35 (Methods). This model had an accuracy in predicting the regime of >95% on the training data from long-standing populations (Figure 4A), whereas on the held-out data from more recently derived populations, the model predicted regime perfectly (Figure 4B). Failing to distinguish the metabolome of recent and long-standing populations along any of the PCs and the high accuracy of discrimination models on the recently derived metabolome suggest that the metabolome has converged by regime among the recently derived populations.

**Figure 3.**
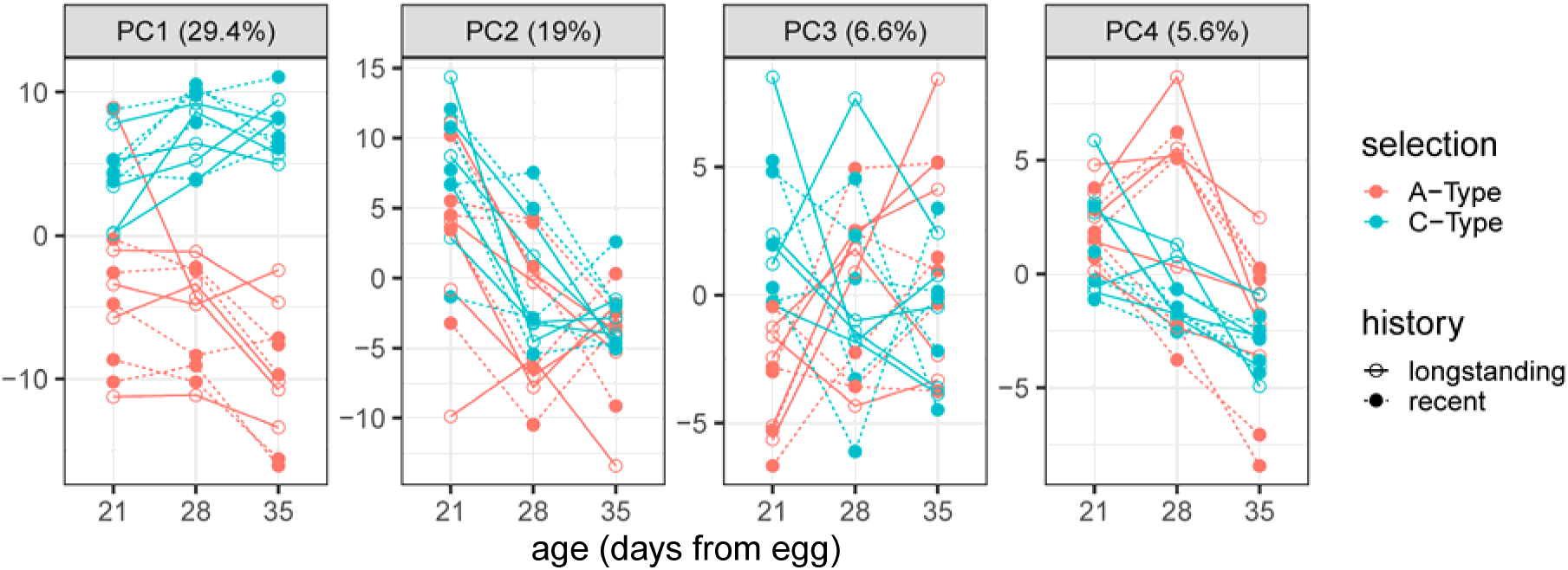
The metabolome and its age-trajectory converge under each selection regime. Each of the first 4 principal components of the metabolome (out of 12 assayed) are plotted over age, from egg (days), for the five biological replicates of each selection regime (A-type or C-type) and selection history (long-standing or recently derived). Ages in which both regimes are sampled are shown for direct comparison. PCs 1 and 4 were highly associated with age by regime, PCs 2 and 4 were associated with age, and PC 3 was associated with regime (ANVOA, P<0.05, after Bonferroni correction). No PCs had a significant effect of history or history x regime interaction which would be inconsistent with convergence of the metabolome by regime in the long-standing and more recently derived populations.

**Figure 4.**
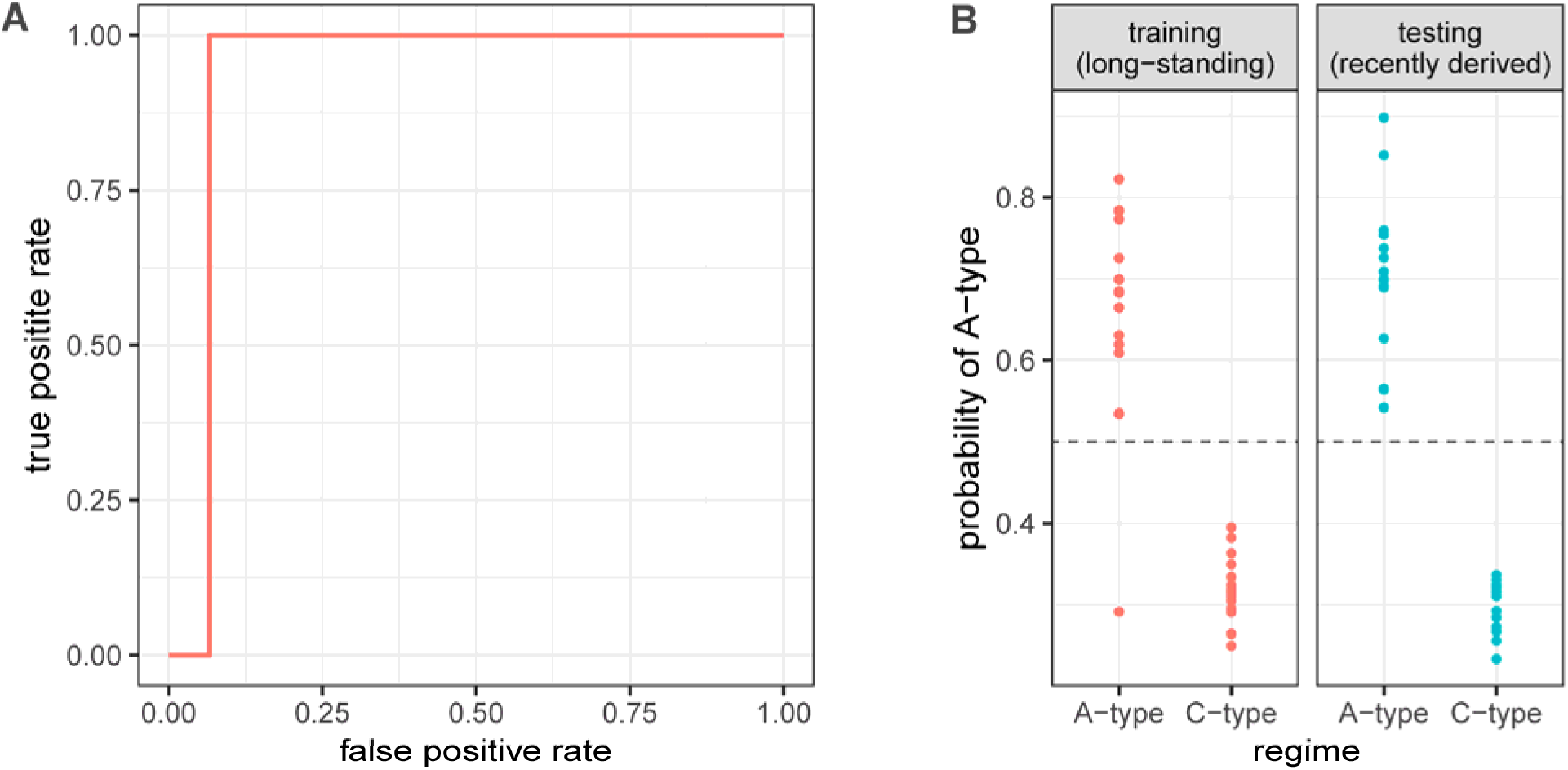
Regime is accurately predicted from the metabolome of the recently derived selection lines. A 2-component partial least-squares discrimination model was trained, by 5- fold cross-validation to distinguish regime (A-type or C-type) among the 30 samples of the long-standing populations using the data from 202 metabolites. A) the ‘ROC’ curve for the training data. Accuracy in the training data was 96.7% in the final model, with an area under the ROC curve of 0.975. B) The probability of A-type for each of the 30 samples from the model training set (training, long-standing, and the 30 withheld samples from the recently derived populations). Discrimination is made at the 0.5 probability (dashed line). The model distinguished all samples in the test set with 100% accuracy.

#### Objective 2: Metabolomic Clocks and the Effect of Selection Regime

Consistent with other phenotypes, we hypothesize that the metabolome will also serve as a predictive phenotype for both chronological and biological age. We expect that, if there is a detectable selection driven alteration to the metabolome, then the A-type populations will appear older relative to the C-type populations of the same chronological age. To test for the signal of aging in the metabolome, we constructed metabolomic “clocks” trained on metabolite abundance patterns specific to each selection regime, then used those clocks to predict the age of samples both within, and between regimes (Methods). To maintain relevant comparisons across all major analyses, data from the 9-day-old A-type samples were withheld when training and testing the clock models due to the unique developmental state of these samples, for which we do not have an equivalent in the C-type samples.

### Metabolomic Clock Results

We first evaluated the relationship between the metabolome and age within the A-type and the C-type populations using within-regime elastic net regression models (clocks, Methods). The within-regime clocks of the C-type and A-type both accurately predicted the age of flies in their respective regimes, with an *R^2^* of 0.87 and 0.77 respectively (Figure 5). To test hypotheses about the differences in biological age, or the pace of aging in response to selection, we turned to between-regime metabolomic clocks (Methods). We fit a linear model to the effect of age on the predicted ages of within and between-regime clocks. Specifically, we test for both age-independent differences in the clock predictions (β*_2_*), which would be expected if selection has led to differences in the apparent age of the metabolome regardless of age, as well as age-dependent differences (β*_3_*), which might reflect differences in the apparent pace of aging in response to selection.

**Figure 5.**
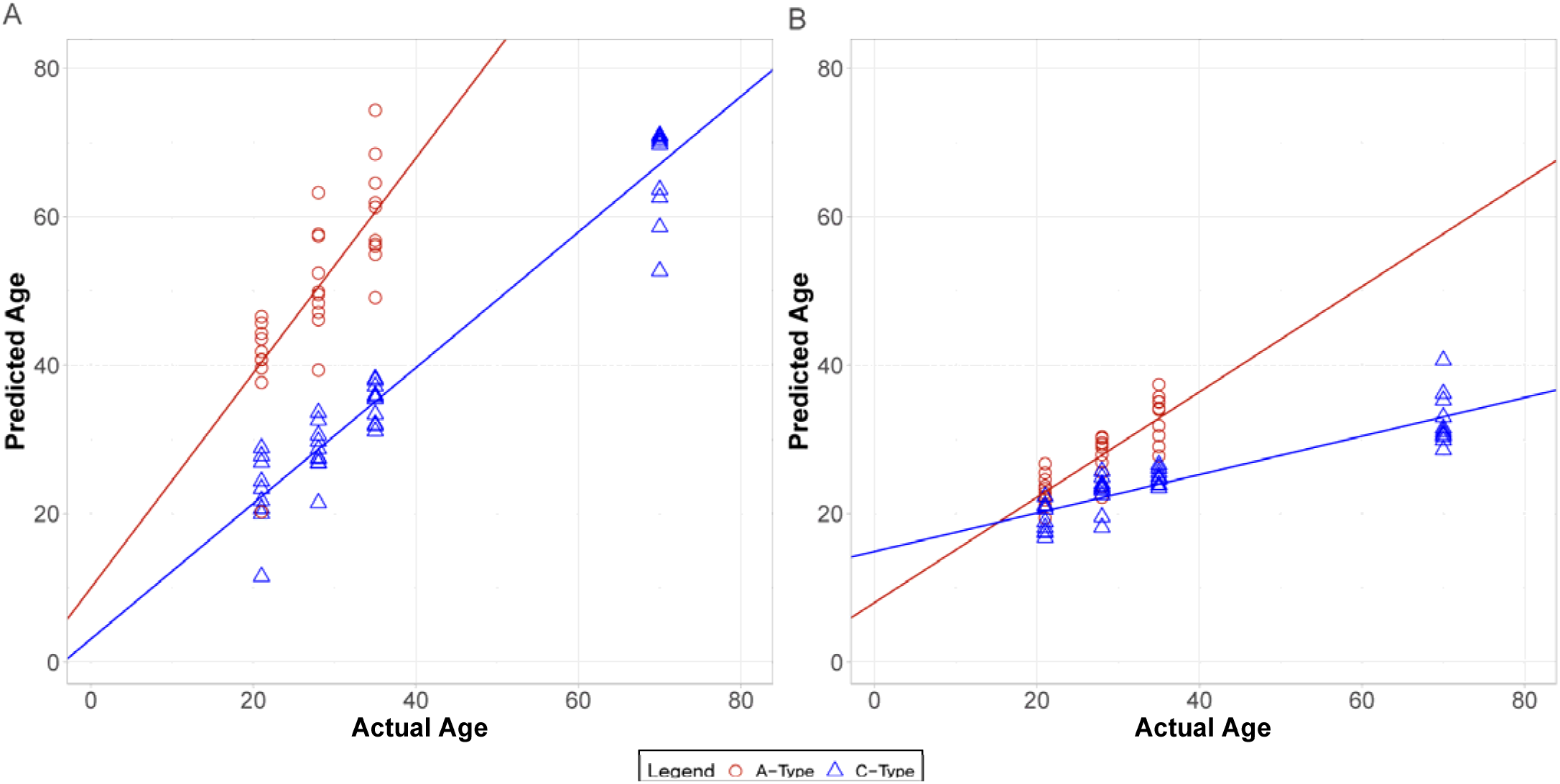
Selection led to different aging trajectories in the metabolome. Predicted age from egg (days) vs chronological age from egg (days) for A-type (red) and C-type (blue) populations based on elastic net models trained on metabolite data from C-type populations (A) or A-type populations (B). Points are predictions from samples held out during training (Methods). The within-regime accuracies of each model were (C-type R^2^ = 0.87; A-type R^2^ = 0.77). Lines are the least-squares fits of a linear model (Methods). Both the A and C-type clocks predicted a significant difference in the rate of aging between the within and between-regime model predictions (A-type clock p = 1.5×10^-6^ and C-type clock p = 9.3×10^-3^), with both predicting a faster rate of aging for A-type flies relative to C-type flies.

We begin by comparing the effect of age on predicted ages made by the C-type clock, when applied to the A-type metabolome, to predictions of the C-type clock on the C-type metabolome (Methods, Figure 5A). The ages predicted from the A-type metabolome by the C-type clock did not differ in intercept (β*_2_*= 6.87, p = 0.25), but had a greater slope than the within-regime predictions of the C-type (β*_3_* = 0.53, p = 9.3×10^-3^). These results suggest that, as estimated by the C-type clock, the A-type metabolome shows a faster rate of aging than expected by its chronological age.

We then used the A-type clock to predict age in C-type flies and the same linear model framework to test for effects of selection regime on aging in the metabolome (Figure 5B). The A-type clock predicted ages from the C-type metabolome that changed more slowly over chronological age than those of the A type (β*_3_* = −0.45, p = 1.5×10^-6^). The y-intercept of predicted age of the C-type was higher than that of the predictions on the A-type (β*_2_* = 6.89, *p* = 8.38×10^-3^). However, the predicted ages of C-type samples from the A-type clock were lower than both their chronological age and the predicted age of A-type samples of the same chronological age (Figure 5B). These results suggest that, as estimated by the A-type clock, the C-type metabolome appears to age at a slower rate than the A-type metabolome.

The A-type and C-type clocks make different age predictions on metabolome data from the two regimes sampled at the same chronological age. And yet, regardless of regime, the two clocks predict increasing biological age as chronological age increases. We considered two possibilities, one, that the two clocks are primarily utilizing variation in the same subset of metabolites within either regime to make age predictions. Alternatively, the A- and C-type clocks could gain predictive power from two different sets of metabolites as they vary with age within-regime. To distinguish these two possibilities, we examined the coefficients fit to each metabolite within both clocks.

Both clocks used an elastic net penalty L1>0, and so at least some metabolites were given no importance. This left 62 metabolites selected as model features for the A-type clock, and 23 metabolites as model features for the C-type clock (Table S4). Of these features, four metabolites were common to both clocks. An intersection of four metabolites does not indicate enrichment of metabolites among the two clocks (Fisher’s test, odds ratio=0.44, P=0.16). However, the coefficients for the four metabolites common to both clocks were highly correlated (Pearson’s ρ=0.97, P=0.035, Figure 6A). In comparison to the importance (β) of the metabolites specific to the C-type or A-type clocks, the four common metabolites were not significantly more or less important within either clock (Wilcoxon’s test W≤155, P≥0.27, Figure 6B).

**Figure 6.**
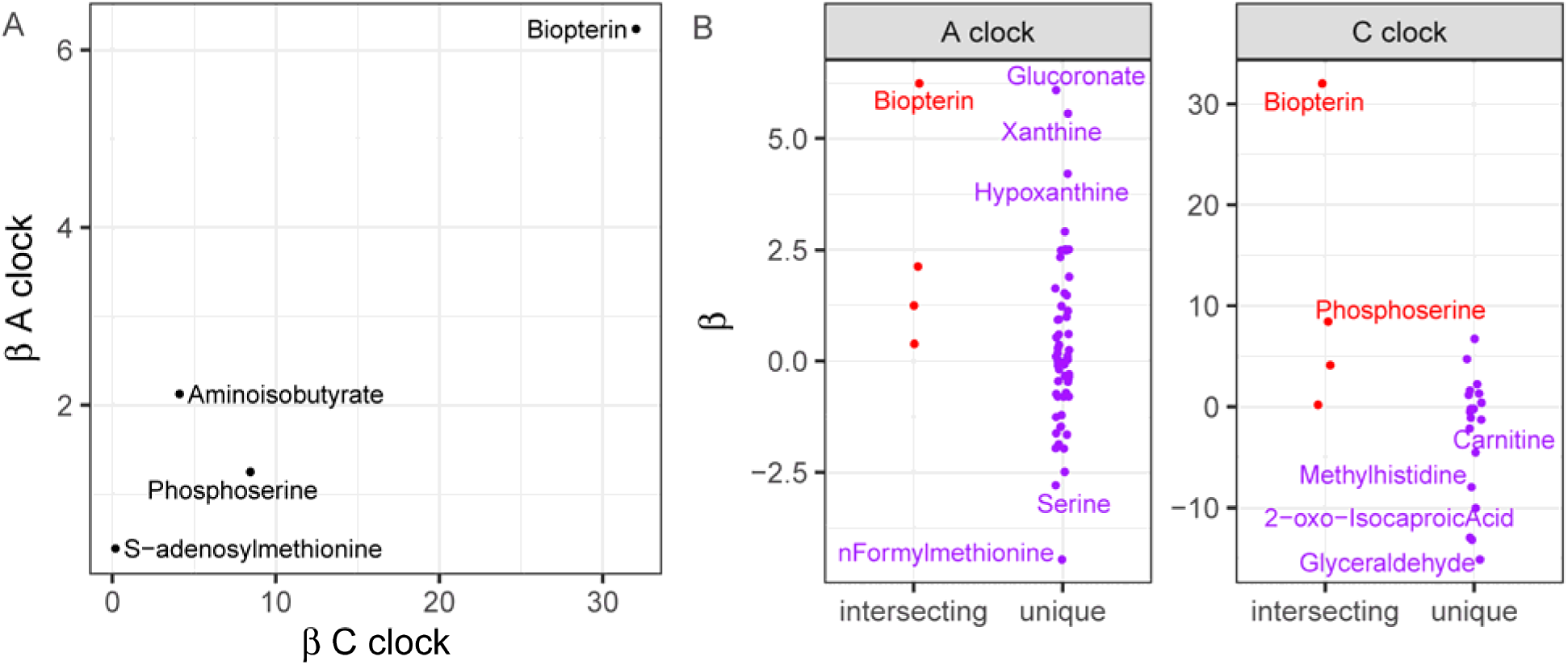
Metabolomic clock features. A: The coefficients (*β*) fit to each of the 4 metabolites common to the A- and C-type clocks (Pearson’s ρ=0.97, P=0.035). B: Comparing the distribution of *β* fit to metabolites common to both clocks (intersecting) and the metabolites unique to each clock. There was not a significant difference in the |*β*| for intersecting and unique metabolites in either clock (Wilcoxon rank sum test, W≤155, P≥0.27).

#### Objective 3: Metabolomic Trajectories of Rapid Aging

To characterize the metabolomic trajectories associated with aging in the populations under the A and C-type selection regimes, we used a multifaceted approach to reveal age and regime specific patterns of metabolite abundance.

First, a linear mixed-effects model (LMM) approach was used to characterize specific patterns of metabolite differentiation (Table S5). Again, we did not include the 9-day-old A-type samples for this analysis due to the unique developmental stage of these samples. Of the 202 metabolites in our data set, a total of 157 unique metabolites were identified as significant for at least one term (FDR <0.01) (Figure 7). The selection and age terms explained most of the difference in the metabolome, accounting for 111 and 112 of the 157 significant metabolites respectively. Both terms consisted of 36 metabolites unique to each term and shared 56 additional metabolites between them. The interaction term accounted for 29 of the significant metabolites, with only 7 metabolites unique to the term.

**Figure 7.**
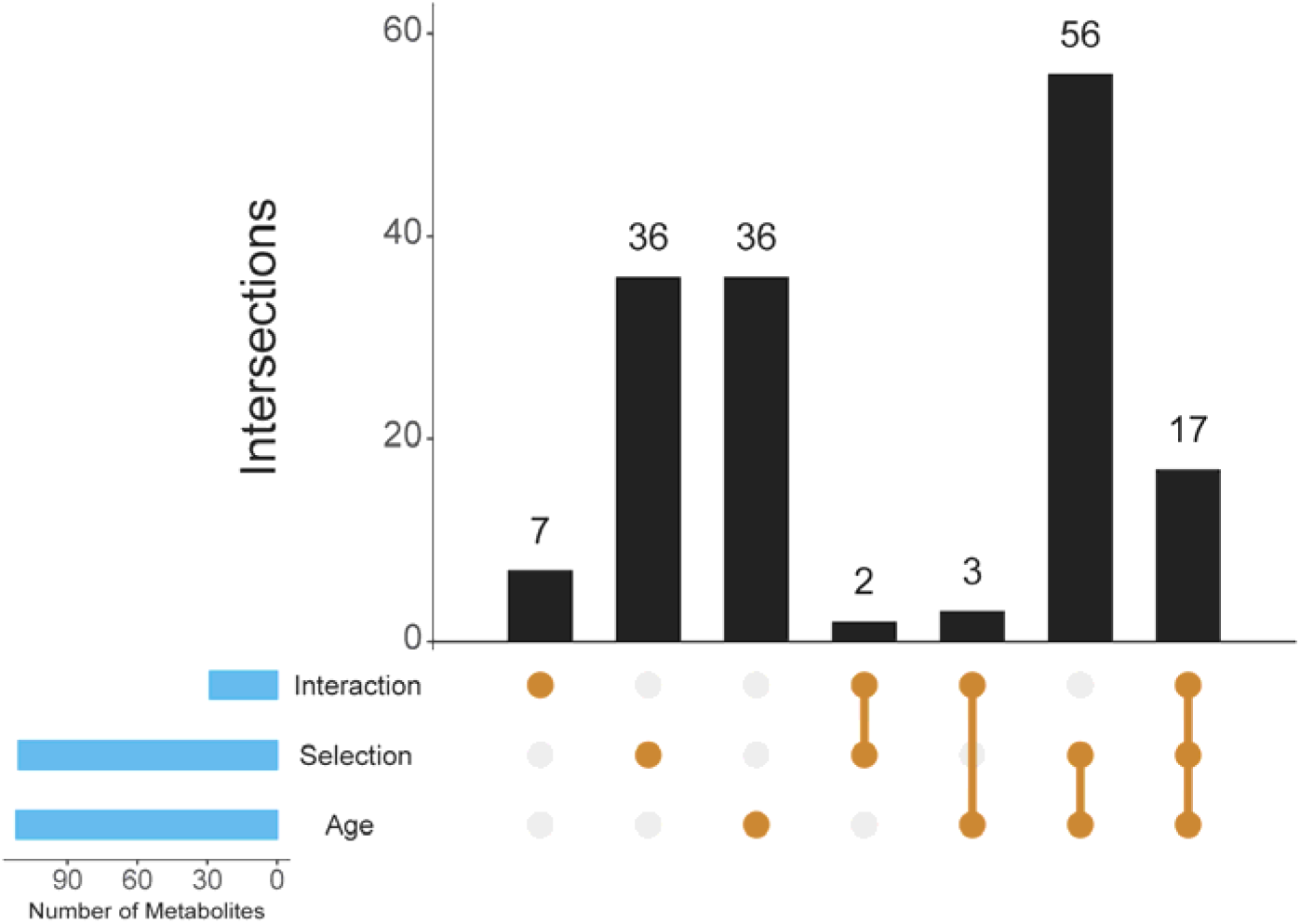
UpSet plot depicting the number of metabolites that were significant from LMM (FDR <0.01) by each term. Selection, age, and the interaction along the lower left, and the number of metabolites that are significant for each term individually and each combination of terms (Intersections) along the top.

### Characterizing Metabolite Abundance Patterns

Next, we generated a heatmap that included the means of normalized values for each age and selection regime for the 202 metabolites detected in all samples (Table S2). Hierarchical clustering based on columns (representing selection regime and age in days) resulted in clusters that are consistent with the earlier PCA (Figure S1), where A-type age 9 samples clustered independently, the matched ages (21, 28, and 35 days) clustered within regime but not across regimes, and the C-type age 70 samples clustered more closely with the matched age samples from the A-type regime than the rest of the C-type samples (Figure S2). Hierarchical clustering of rows (mean metabolite abundance) revealed 25 clusters, five of which exhibited a pattern where A-type samples (aged 21, 28, 35 days) and the oldest C-type samples (aged 70 days) have metabolite abundance patterns more similar to each other than to the younger C-type samples. This pattern is consistent with the general accelerated aging phenotype observed in the A-type populations, potentially indicating an “aged phenotype” represented in the metabolome. To highlight this pattern, we used the subset of metabolites included in these five clusters to generate a set of reduced heatmaps, where column clustering was maintained from the full heatmap, and rows were allowed to cluster based on the subset data, both resulting in dendrograms consistent with the larger heatmap (Figure 8). These clusters generally represent either metabolites with higher abundance in the samples with an aged phenotype (clusters 6 and 10), or lower abundance in samples with an aged phenotype (clusters 2, 9, and 22) relative to the younger C-type samples. These clusters contained a total of 68 of the 202 metabolites assayed (Full list of metabolites by cluster can be found in Figure S2). When compared to the LMM results, all clusters contained at least one metabolite that was significant (FDR < 0.01) for at least one term from the LMM (either selection, age, or the interaction between the two), representing an average of 85.3 % (range 25 – 100 %) of the metabolites in each cluster significant for at least one term (Table 1).

**Figure 8.**
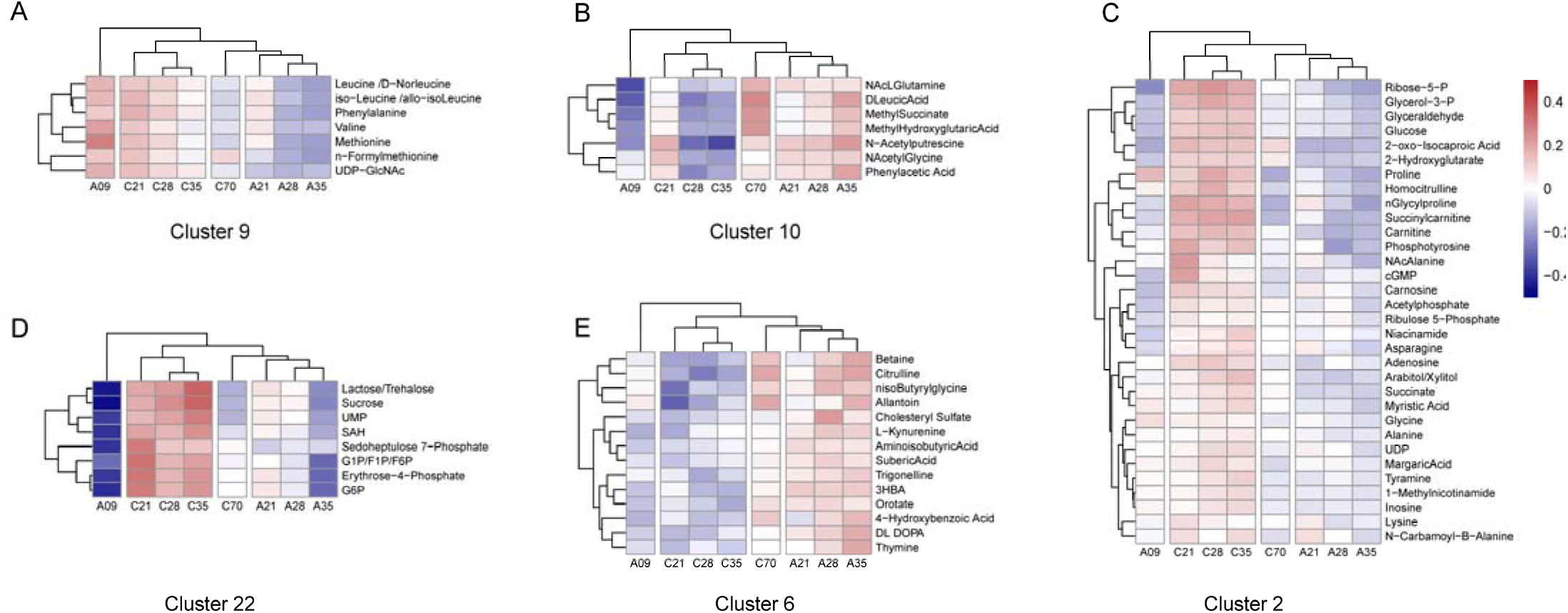
Hierarchal clustering of mean normalized metabolite abundance clustered by regime and age. The dendrograms were generated using 202 metabolites that were detected in all samples (see Figure S2 for heatmap that includes all 202 metabolites). These five clusters represent examples where C-type age 70 days have metabolite abundance that more closely resembles the A-types than the C-types at younger ages. KEGG network analysis revealed enrichment for several pathways for each cluster. The top pathway for each cluster, representing the lowest p-score among significant pathways for that cluster (FDR<0.05), included A: Valine, leucine and isoleucine biosynthesis (dme00290); B: Phenylalanine metabolism (dme00360); C: Pentose and glucuronate interconversions (dme00040); and D: Glycolysis / Gluconeogenesis (dme00010); E: Nitrogen metabolism (dme00910).

**Table 1.**
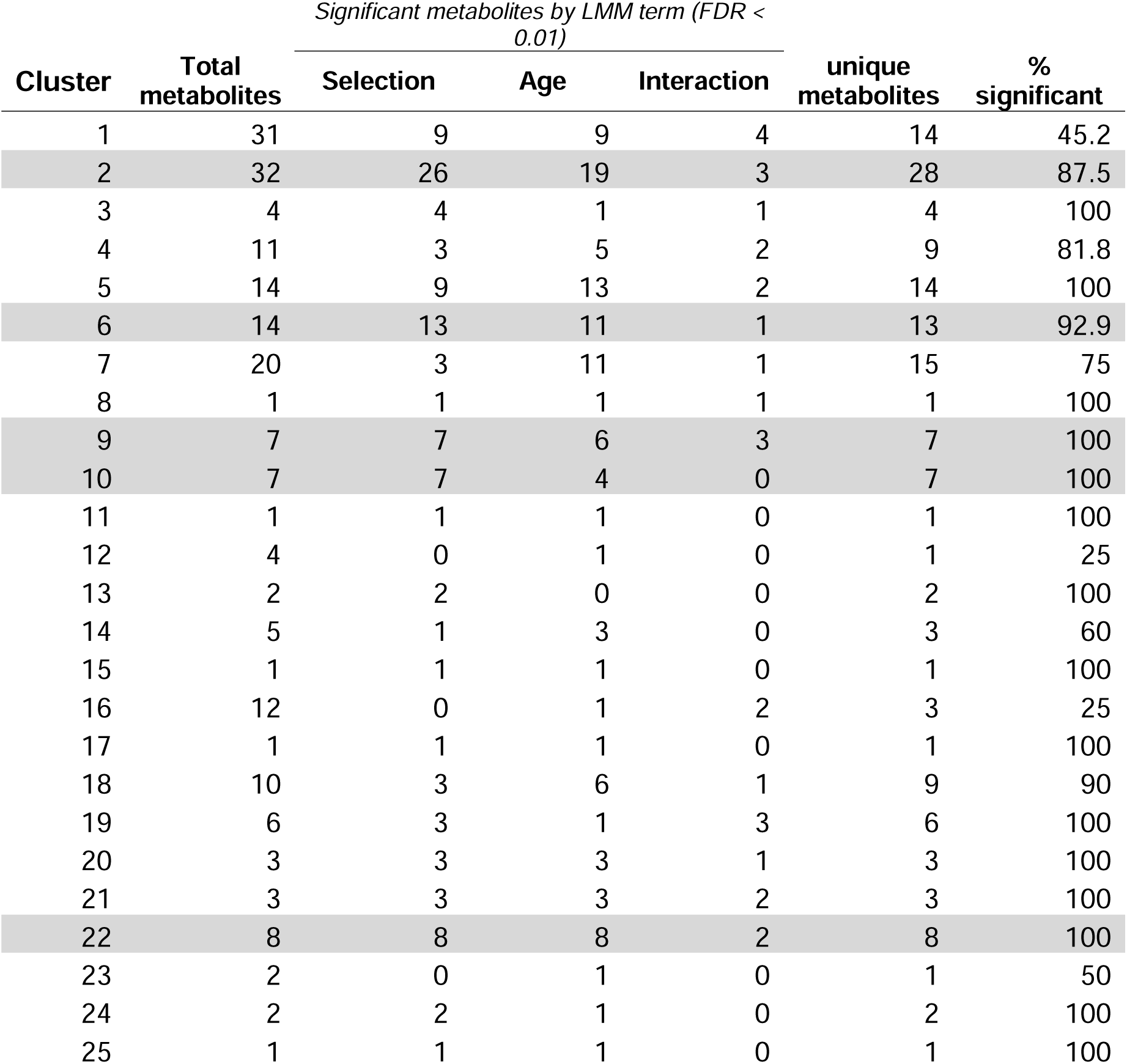
Breakdown of metabolites by hierarchical clustering of mean normalized metabolite abundance and significant results from LMM (Table S5). Shaded rows indicate clusters that show abundance patterns consistent with the accelerated aging phenotype (Figure 8).

| Cluster | Total metabolites | Significant metabolites by LMM term (FDR < 0.01) |  |  | unique metabolites | % significant |
| --- | --- | --- | --- | --- | --- | --- |
|  |  | Selection | Age | Interaction |  |  |
| 1 | 31 | 9 | 9 | 4 | 14 | 45.2 |
| 2 | 32 | 26 | 19 | 3 | 28 | 87.5 |
| 3 | 4 | 4 | 1 | 1 | 4 | 100 |
| 4 | 11 | 3 | 5 | 2 | 9 | 81.8 |
| 5 | 14 | 9 | 13 | 2 | 14 | 100 |
| 6 | 14 | 13 | 11 | 1 | 13 | 92.9 |
| 7 | 20 | 3 | 11 | 1 | 15 | 75 |
| 8 | 1 | 1 | 1 | 1 | 1 | 100 |
| 9 | 7 | 7 | 6 | 3 | 7 | 100 |
| 10 | 7 | 7 | 4 | 0 | 7 | 100 |
| 11 | 1 | 1 | 1 | 0 | 1 | 100 |
| 12 | 4 | 0 | 1 | 0 | 1 | 25 |
| 13 | 2 | 2 | 0 | 0 | 2 | 100 |
| 14 | 5 | 1 | 3 | 0 | 3 | 60 |
| 15 | 1 | 1 | 1 | 0 | 1 | 100 |
| 16 | 12 | 0 | 1 | 2 | 3 | 25 |
| 17 | 1 | 1 | 1 | 0 | 1 | 100 |
| 18 | 10 | 3 | 6 | 1 | 9 | 90 |
| 19 | 6 | 3 | 1 | 3 | 6 | 100 |
| 20 | 3 | 3 | 3 | 1 | 3 | 100 |
| 21 | 3 | 3 | 3 | 2 | 3 | 100 |
| 22 | 8 | 8 | 8 | 2 | 8 | 100 |
| 23 | 2 | 0 | 1 | 0 | 1 | 50 |
| 24 | 2 | 2 | 1 | 0 | 2 | 100 |
| 25 | 1 | 1 | 1 | 0 | 1 | 100 |

Finally, cluster-specific functional enrichment analysis (FELLA) revealed 86 enriched pathways (FDR < 0.05) across the 15 clusters that contained at least four metabolites with KEGG annotation (Table S6). Among the five clusters representing the accelerated aging phenotype, 21 enriched pathways were discovered (Table 2). These 21 enriched pathways were involved primarily in carbohydrate metabolism (clusters 2 and 22), nitrogen metabolism (clusters 6 and 10), and growth and development (clusters 6 and 9), suggesting that these metabolites may play an underlying role in the different rates of development and aging we see in this system.

**Table 2.** Enriched pathways for clusters of metabolites with abundance patterns consistent with the accelerated aging phenotype (Figure 8). Shaded cells represent clusters (6 & 10) with higher metabolite abundance in the aged phenotype, and unshaded cells (2, 9, & 22) represent clusters with lower metabolite abundance in the aged phenotype. See Table S6 for full list of significant KEGG pathways and modules.

| Cluster | KEGG ID | KEGG name | p.score | FDR |
| --- | --- | --- | --- | --- |
| 2 | dme00040 | Pentose and glucuronate interconversions | 0.0221 | 0.0497 |
|  | dme00760 | Nicotinate and nicotinamide metabolism | 0.0267 | 0.0497 |
|  | dme00780 | Biotin metabolism | 0.0321 | 0.0497 |
|  | dme00030 | Pentose phosphate pathway | 0.0366 | 0.0497 |
| 6 | dme00910 | Nitrogen metabolism | 0.0018 | 0.0049 |
|  | dme00100 | Steroid biosynthesis | 0.0023 | 0.0051 |
|  | dme00981 | Insect hormone biosynthesis | 0.0038 | 0.0056 |
| 9 | dme00970 | Aminoacyl-tRNA biosynthesis | 0.0006 | 0.0147 |
|  | dme00290 | Valine, leucine and isoleucine biosynthesis | 0.0017 | 0.0163 |
|  | dme04980 | Cobalamin transport and metabolism | 0.0140 | 0.0238 |
|  | dme04330 | Notch signaling pathway | 0.0152 | 0.0253 |
|  | dme03083 | Polycomb repressive complex | 0.0204 | 0.0274 |
| 10 | dme00360 | Phenylalanine metabolism | 0.0007 | 0.0062 |
|  | dme00982 | Drug metabolism - cytochrome P450 | 0.0094 | 0.0144 |
| 22 | dme00010 | Glycolysis / Gluconeogenesis | 0.0001 | 0.0003 |
|  | dme00030 | Pentose phosphate pathway | 0.0001 | 0.0003 |
|  | dme00052 | Galactose metabolism | 0.0001 | 0.0003 |
|  | dme03018 | RNA degradation | 0.0001 | 0.0003 |
|  | dme00500 | Starch and sucrose metabolism | 0.0002 | 0.0004 |
|  | dme00604 | Glycosphingolipid biosynthesis - ganglio series | 0.0013 | 0.0016 |
|  | dme00051 | Fructose and mannose metabolism | 0.0016 | 0.0016 |

## Discussion

### Objective 1: Metabolomic Convergence Within Selection Regime

Adaptation in response to selection on reproductive timing in this system is associated with a number of physiological and developmental tradeoffs with complex genetic underpinnings. Because the metabolome constitutes the functional building blocks, and energetic currency of cells, we hypothesize that the physiological tradeoffs associated with the accelerated aging phenotype of A-type flies would be reflected in the abundances of metabolites. Alternatively, the metabolome could be associated with a wide variety of traits, only some of which are involved in the accelerated development or reduced longevity that evolve under these regimes. If our hypothesis is true, then we expect that populations under the same selection regime will show evidence of convergence within the metabolome, even in more recently selected populations. To test this prediction, we compared the metabolomes of populations from recently derived and long-standing histories under the same selection regime. We found that the metabolomes of recently derived and longstanding populations did not differ within either the A- and C-type selection regimes. This was evident by both a lack of metabolomic divergence between long standing and more recent selection within each regime, and by the high accuracy of multivariate model predictions, regardless of the number of generations under selection. These results are consistent with metabolomic convergence within selection regimes and indicate that the metabolome is a reproducible biomarker of the underlying physiological responses to selection.

### Objective 2: Metabolomic Clocks and the Effect of Selection Regime

The dynamics of the metabolome over the lifespan of the fly have been used to connect advancing chronological age to the rise in mortality risk that defines biological aging (Zhao et al. 2022). Multivariate age prediction models (clocks), particularly epigenetic clocks, have been used to predict age across species (Meyer and Schumacher 2024; Lu et al. 2023). The use of clock models to understand the evolutionary process however is just beginning (Parrott and Bertucci 2019). Here we apply clock models of the metabolome to test our hypothesis that the rapid response to selection for early life reproduction is associated with the same underlying aging processes that are reflected in the metabolome. If true, the response to selection seen in the metabolome likely reflects shifts in physiological processes that are a part of normal aging in the ancestral population. If the rapid and repeatable shifts in allele frequency (Graves et al. 2017), and both transcriptomic (Barter et al. 2019) and phenotypic (Burke et al. 2016; Kezos et al. 2023) convergence are an indication of selection acting on common physiological mechanisms, then the components of the metabolome most closely tied to aging physiology should respond similarly. A critical prediction of our hypothesis is that the relative biological age of flies under the A-type selection regime should appear ‘older’ than flies under the C-type selection regime.

To test this hypothesis, we first trained age prediction models *within* each regime, resulting in an A-type clock and C-type clock. Both clocks captured the monotonic increase in chronological age within each respective population, with accuracy R^2^ exceeding >0.77 (Figure 5). In between-regime predictions, the C-type clock detected a chronological age increase for the A-type flies over time, however these age predictions were overestimated across all ages. Additionally, the C-type clock predicted an increase in the rate of aging of A-type flies relative to the predicted aging rate of flies form the C-type regime (Figure 5A). From a metabolomic perspective, the C-type flies “see” the A-type flies as both older and aging more quickly, presenting a metabolomic signal of accelerated biological aging in flies under the A-type selection regime.

Between-regime predictions using the A-type clock detected a chronological age increase for the C-type flies over time, however this prediction was underestimated across all ages. Additionally, the A-type clock predicted a decrease in the rate of aging of C-type flies relative to the predicted aging rate of flies from the A-type regime (Figure 5B). From a metabolomic perspective, the A-type flies “see” the C-type flies as both younger and aging more slowly, presenting a metabolomic signal of a slower biological aging in flies under the C-type selection regime.

Taken together, the differences in age predictions made by the within-regime clocks show that the metabolome is capturing a biological signal that is indicative of chronological age, while the between-regime clocks show that the metabolome is also able to capture a signal indicative of the differences in biological age and aging observed at other phenotypic levels (e.g. Chippindale et al. 1997; Kezos et al. 2023; Burke et al. 2016). The A-type clock suggests an age-deceleration effect when applied to the metabolome of the longer-lived C-type populations and the C-type clock suggests an age-acceleration effect when applied to the shorter-lived A-type populations. Indeed, even the effects on the slopes are remarkably similar between the two between-regime models, both featuring a difference in a 50% difference in slopes. This further suggests both consistency and reliability in the metabolome for the process of aging even across population with different lifespans.

The A-type clock achieved an accuracy of R^2^=0.77 when predicting age in the A-type, while utilizing only 30.7% of 202 metabolites, and the C-type clock utilized only 11.4% to achieve even higher accuracy R^2^=0.87. Independently fit to two different data, the C-type and A-type clocks utilized predominantly non-intersecting metabolites, however they both included four intersecting metabolites (Figure 6A, Table S4). While this number of intersecting metabolites is not more than expected by chance, the coefficients fit to these metabolites in both clocks were highly correlated, indicating a common axis of variation with age in the metabolome.

It is important to note that, while the metabolites identified by the clock models shed light on the underlying metabolome variation related to age and to aging, elastic net and other regularized regression methods are designed in part to reduce the influence of co-linear predictors (Zou and Hastie 2005; Bujak et al. 2016). Because of this, the set of metabolites most influential in clock models may not include metabolites that share very similar age-related variation in abundance and so do not give a complete picture of age-associated metabolome variation in which collinearity is common (Lassen et al. 2023; Avanesov et al. 2014). In this light, we turn to a descriptive analysis of the metabolome variation and its convergence within regime to describe the patterns of variation under these selection regimes.

### Objective 3: Characterizing Metabolite Abundance Patterns

Before we delve deeper into pathway analysis, we must first address the overall patterns of relative metabolite abundances across age and between populations under our two selection regimes. When evaluating the results of the linear mixed-effects models, it is important to take special note of the influence from the age x regime interaction term, as it helps to establish perspective for subsequent analysis. Based on the results from the PCA (Figure 3) and the clock model predictions (Figure 5) we see evidence of a large effect from the selection regime on the metabolome, as predicted age, and rate of aging differ by regime. As such, this led us to expect to see this reflected in the LMM results as a high number of metabolites that are significant for the age x regime interaction term, where many metabolites are contributing the accelerated aging phenotype of the A-type populations. However, we were surprised to find the opposite to be true, where relatively few metabolites were significant for the age x regime interaction term (14% of total metabolites) compared to the selection and age terms (55% each), suggesting that only a handful of metabolites may be driving the accelerated aging phenotype. With this in mind, we move forward with hierarchical clustering of metabolite abundance and pathway analysis to derive further insight into how small clusters of metabolites might drive these broader aging phenotypes.

To discuss the implications of the differences in metabolite abundance between selection regimes and across age, we must make assumptions about how metabolite abundances relate to the activity of specific metabolic pathways. For example, we could assume that increased abundance of a metabolite indicates increased activity of the pathway that produces it. However, as metabolites are often involved in multiple pathways – both as substrate and product – reality is likely more complex. Other factors like differences in age-specific feeding rates can also impact abundances in ways that confound interpretation of abundances. We attempt to overcome this, in part, through the use of higher-level analyses such as hierarchical clustering and enrichment analysis instead of making direct interpretations of individual metabolite abundances. As such, we primarily see the following work as hypothesis generating for future studies (e.g., flux assays to test hypotheses about metabolite utilization and pathway activity).

The metabolite abundance patterns driving the clusters depicted in Figure 8 are consistent with the prediction that the older C-type (70-day-old) and younger A-type (21, 28, and 35-day-old) flies would share similar patterns of abundance, revealing a metabolomic phenotype indicative of advanced or accelerated biological age (what we will refer to as the “aged phenotype”). This similarity in the metabolomes of old C-type and younger A-type flies is consistent with a broad array of age-related phenotypes shared by these two groups of flies, including high mortality rates (Figure 2), as well as reduced egg viability (Chippindale et al. 1997), stress resistance (Kezos et al. 2023), and lifetime fecundity (Burke et al. 2016). These clusters represent two distinct patterns that may help shed light on the mechanisms underlying the accelerated aging phenotype of flies under the A-type selection regime.

The first pattern, exhibited in clusters 9, 2, and 22 (Figure 8A, C, and D, respectively), include metabolites with a lower abundance in flies with the aged phenotype, suggesting that pathways associated with these metabolites may be utilized less in these biologically older flies. Clusters 2 and 22 both show enrichment for several pathways involved in carbohydrate metabolism, most notably the decrease in metabolites enriching the glycolysis and gluconeogenesis pathway in cluster 22 (Table 2). This decreased abundance of metabolites involved in carbohydrate metabolism suggests a potential switch in metabolic substrates away from carbohydrates that is associated with the aged phenotype (Hoffman et al. 2014; Wang et al. 2022). Along with the decrease in carbohydrate metabolism, we also see a decreased abundance of metabolites associated with protein synthesis enriched in cluster 9, such as enrichment for aminoacyl-tRNA biosynthesis and the valine, leucine and isoleucine biosynthesis pathway. The pattern of metabolite abundance in clusters 10 and 6 (Figure 8B and E respectively) include metabolites with increased abundance in flies with the aged phenotype suggesting that pathways associated with these clusters may be utilized more in these biologically older flies. The top pathways enriched for these clusters include both phenylalanine and nitrogen metabolism (cluster 10 and 6, respectively; Table 2), both of which are employed in circumstances with low carbohydrate utilization (Heinrichsen et al. 2014; Kosakamoto et al. 2022).

In considering the decrease in metabolite abundances associated with glycolysis, gluconeogenesis, and protein synthesis pathways, alongside the increase in metabolite abundances associated with nitrogen metabolism pathways, we propose this is evidence of a shift away from carbohydrate metabolism towards amino acid metabolism in flies with the aged phenotype. This pattern of age-related energy substrate use has been previously reported in *Drosophila*, where Wang et al. (2022) reported a decrease in carbohydrate use and an emphasis on serine and purine metabolism in aged flies. Additionally, Phillips et al. (2022) reported a reduction in both amino acid and carbohydrate abundances in 21-day-old A-type flies relative to C-type flies. Thus, the apparent shift we see reflected over the time-course measured here could explain the initial observations of Phillips et al., 2022. While more work is needed to directly test for such a substrate switch, it seems likely that the earlier age of onset of this metabolomic change in A-types is driven by selection for early reproduction. The accelerated development and heightened acute fecundity in flies from the A-type selection regime (Burke et al. 2016) may be driving an antagonistic pleiotropic effect, where selection for carbohydrate utilization strategies that support rapid development and oviposition comes at the cost of an earlier shift away from this metabolic pathway, typically only seen in much older individuals, potentially contributing to the accelerated aging phenotype (Hoffman et al. 2014; Wang et al. 2022).

## Materials and Methods

### Experimental Populations

All 20 *D. melanogaster* populations used in this study share a common origin, traced back to the large and outbred Ives (1970) population. Numerous populations have been derived from this wild-bred lineage, first by Michael Rose and Brian Charlesworth (Charlesworth 1980; Rose & Charlesworth 1980; Rose & Charlesworth 1981; Rose 1984) and then by generations of the Rose Lab (e.g. Chippindale 1997, Burke et al. 2016), to study the evolutionary theory of aging. All populations in this system are maintained at census sizes of ∼2000 individuals in an attempt to keep them outbred. Based on past genomic studies, we know there is abundant genetic variation in all populations even after many hundreds of generations of evolution (e.g. Phillips et al. 2016; Graves et al. 2017). Here we focused on the A-type populations and their controls, the C-type populations (Figure 1B). A-type populations were maintained on a 10-day generation cycle, whereas the C-type populations were maintained on a 28-day cycle. As a result of this difference, the A-type populations exhibit accelerated development, reduced longevity, higher early-life fecundity, and lower stress tolerances than to the C-type populations (Burke et al. 2016; Kezos et al. 2023). There are two selection histories within the A and C-type populations, ACO/AO and CO/nCO, respectively (See Figure 1A). Each history is distinguished by the number of generations they have been under selection, with AO and nCO having been derived more recently than ACO and CO to test questions about evolutionary repeatability (see Burke et al. 2016 and Graves et al. 2017). We know that populations from these selection histories that share the same selection regime have converged at several phenotypic levels and recent studies have treated them as replicates within each regime (A or C), based on phenotypic (Burke et al. 2016), genomic (Graves et al. 2017), and transcriptomic (Barter et al. 2019) studies. At the time of sampling for this project, the ACO and AO populations had ∼1,000 and ∼490 generations under selection, respectively, whereas the CO and nCO populations had ∼415 and ∼160 generations, respectively. In all of the experiments described here, we include five historical replicate populations of each of the four experimental populations (ACO, AO, CO and nCO), that have been maintained as independent lineages for the entirety of the generations under each selection regime.

### Mortality Assays

Adult mortality assays were based on published protocols (Burke et al., 2016) which measured mortality in the same five replicate populations of each of the four experimental populations ACO, AO, CO, and nCO. Prior to measurement, all of the experimental populations were maintained on a 14-day culture cycle for two generations, staggered a day apart, to align the different treatments to a shared calendar and reduce the impact of gene by environment interaction. Each replicate was expanded into two cohorts (e.g. ACO1α & ACO1β) of ∼1,500 flies, each in a Plexiglas cage, supplied with fresh banana-molasses based media (as described in Phillips et al. 2018) each day with yeast solution to induce oviposition once transferred on day 14 from egg. Ambient temperature was maintained at ∼25 °C throughout. At the same time each day, all dead flies were removed from each cage to be sexed and logged until no living flies remained. Mortality data was pooled between the two cohorts for each population to give the total number of deaths per day of each population. Age specific mortality (*M*) was calculated as follows:

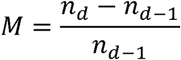

Where *n* equals the number of flies alive on a given day (*d*), and *d* −1 represents the day prior. The Burke et al. (2016) protocol was modified for this experiment to remove the use of carbon dioxide in condensing cohorts when census densities were sufficiently low. Instead, flies within cohorts remained in the same cage for the duration of the experiment, regardless of census density, and the inside of the cages were cleaned as needed to remove waste build-up.

### Sample Collection

Samples were collected for metabolomics using protocols described by Phillips et al. (2022). A cage-cohort of ∼1,500 flies was generated from each of the five replicate populations for ACO, AO, CO, and nCO and maintained on a 14-day culture cycle for two generations, staggered a day apart to mitigate the impact of gene-by-environment interactions. Cohorts were maintained in a Plexiglas cage and were fed fresh banana-molasses based media each day with yeast solution to induce oviposition. Concurrent to the mortality experiment, the five replicate cage-cohorts for each selection history were sampled for metabolomic extraction at multiple timepoints (9 days for A-type populations, 21, 28, and 35 days for all populations, and 70 days for C-type populations). Metabolomic profiles are highly sexually dimorphic within the *Drosophila* system (Hoffman et al. 2014). In order to facilitate the large sample sizes and sampling time points needed, as well as to reduce the potential noise caused by using both sexes within pooled samples, we focused on females in our sampling effort. Each sample consisted of a pool of ∼50 female flies that were randomly drawn from each population per timepoint. An equivalent number of males were removed from populations for each sample to maintain population sex ratios. While metabolomic characterization can be done with far fewer than 50 flies, the populations featured in this study are outbred and still harbor a great deal of genetic variation (see Graves et al. 2017). Using pool sizes of ∼50 flies was an attempt to capture and represent this variation. The pooled samples were immediately dry frozen in liquid nitrogen and stored at −80°C prior to metabolite extraction. Due to decreased population sizes at late ages, two samples contained less than 50 flies: ACO_1_ day 35 (n= 25), and nCO_4_ day 70 (n= 36). These populations were selected based on their generation cycle rather than the number of days since eclosion, so age is consistently referred to as the time since egg deposition throughout the manuscript.

The time points used in this study were informed by demographic data and prior studies of flies under these regimes. Burke et al. (2016), and Barter et. al (2019) characterized mortality rates within the A-type and C-type regimes and they defined the “aging” or “non-aging” phases of their lifespan. For the A-type regime, day 9 from oviposition represents a uniquely non-aging phase of the accelerated lines whereas day 21 shows the onset of a rapid aging phase (Figure 2). Conversely, the C-type regimes were sampled in parallel on days 21, 28, and 35, but were only expected to exhibit the aging phase starting at day 35 (Figure 2).

### Sample Preparation

Aqueous metabolites for targeted liquid chromatography–mass spectrometry (LC-MS) profiling of 80 fly samples were extracted using previously described protein precipitation method (Kurup et al. 2021; Meador et al. 2020). Briefly, samples were homogenized in 200 µL purified deionized water at 4 °C, and then 800 µL of cold methanol containing 124 µM 6C13- glucose and 25.9 µM 2C13-glutamate was added. Internal reference standards were added to the samples to monitor sample preparation. Next, samples were vortexed, incubated for 30 minutes at −20°C, sonicated in an ice bath for 10 minutes, centrifuged for 15 minutes at 14,000 rpm at 4°C, and then 600 µL of supernatant was collected from each sample. Lastly, recovered supernatants were dried on a SpeedVac and reconstituted in 0.5 mL of LC-matching solvent containing 17.8 µM 2C13-tyrosine and 39.2 3C13-lactate, and internal reference standards were added to the reconstituting solvent to monitor LC-MS performance. Samples were transferred into LC vials and placed into a temperature-controlled autosampler for LC-MS analysis.

### LC-MS Assay

Targeted LC-MS metabolite analysis was performed on a duplex-LC-MS system composed of two Shimadzu UPLC pumps, CTC Analytics PAL HTC-xt temperature-controlled auto-sampler and AB Sciex 6500+ Triple Quadrupole MS equipped with ESI ionization source (Meador et al. 2020). UPLC pumps were connected to the autosampler in parallel and were able to perform two chromatography separations independently from each other. Each sample was injected twice on two identical analytical columns (Waters XBridge BEH Amide XP) performing separations in hydrophilic interaction liquid chromatography mode. While one column was performing separation and MS data acquisition in ESI+ ionization mode, the other column was equilibrated for sample injection, chromatography separation and MS data acquisition in ESI- mode. Each chromatography separation was 18 minutes (total analysis time per sample was 36 minutes), and MS data acquisition was performed in multiple-reaction-monitoring mode. The LC-MS system was controlled using AB Sciex Analyst 1.6.3 software. The LC-MS assay targeted 361 metabolites and 4 spiked internal reference standards. Measured MS peaks were integrated using AB Sciex MultiQuant 3.0.3 software. Up to 210 metabolites and 4 spiked standards were measured across the study set, and over 90% of measured metabolites were measured across all the samples. In addition to the study samples, two sets of quality control (QC) samples were used to monitor the assay performance and data reproducibility. One QC [QC(I)] was a pooled human serum sample used to monitor system performance, and the other QC [QC(S)] was pooled study samples, were used to monitor data reproducibility. Each QC sample was injected for every 10 study samples. We assessed the reproducibility of the LC-MS using the coefficient of variation (CV) of each of the 202 metabolites among 10 technical replicates. The mean CV was 0.0044 with a range of 6×10^-4^ to 0.029 (Figure S3). Raw results from LC-MS can be found in Table S1.

### Data Processing and Normalization

The LC-MS data was filtered and normalized prior to analysis. Any metabolites with missing values were removed from the data set, and relative peak areas for the remaining 202 metabolites were log-transformed to approximate a Gaussian distribution. Next, within sample metabolite data were mean-centered to account for sample-to-sample variation. Given that the LC-MS profiling was performed in three separate batches over three days, we estimated the main effects of batch (*B*) on each metabolite and corrected for these batch effects by taking the residuals (ε) of the following linear model:

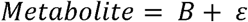

These residuals then constitute the normalized abundance measures for 202 metabolites across 80 fly samples (Table S2).

#### Objective 1: Metabolomic Convergence Within Selection Regime

To test for divergence between recent and long-standing populations in the multivariate metabolome, we first conducted a Principal Components (PC) Analysis (PCA) on the covariance matrix of normalized metabolite abundance using the prcomp R package. We then fit the fixed effects of age (as an ordered categorical factor), regime (A-type or C-type), history (either recent or long-standing), their interactions, and a random effect of replicate on the eigenvalues of each of the first 12 PCs (*PC_X_*) by ordinary least squares in the lme4 R package (Bates et al. 2015).

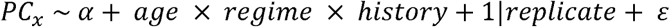

This analysis was limited to the most directly comparable data between regimes, those from day 21, 28, and 35. As a liberal test for divergence, a result that is contrary to our hypothesis, we simply ask if there was either a regime x history interaction effect, *or* a regime x history x age effect on *any* of the first 12 PCs at a P value of <0.05, without adjusting for the 12 tests (each of PC 1 to 12).

To test the potential of the metabolome of long-standing populations to predict selection regime in the recently derived populations, we trained a partial least-squares model to predict regime with the data from the long-standing populations, by 5-fold cross-validation (CV) in the caret R package. This model was then used to predict regime among the samples of the more recent populations. In both training and prediction, only the data from day 21, 28, and 35 were used. The accuracies of the model on the training data (long-standing), and on the held-out data from the recent populations was assessed by receiver operating characteristic (ROC) curve.

#### Objective 2: Metabolomic Clocks and the Effect of Selection Regime

##### Elastic Net Regression Models and Analysis

We used two approaches to generate predicted ages for the flies based on the metabolome data, both of which needed a different approach to, in all cases, avoid age predictions for biological replicates that were part of the model training data. The first approach, whose goal was to generate *within-regime* predictions, required clocks trained and then tested on data from the same regime. The second approach, whose goal was to generate regime-specific models only to then predict ages for the opposite regime. These later clocks we refer to as *between-regime*. The day 9 A-type samples were withheld from these analyses, as results from the PCA (Figure S1) indicate a difference in the metabolomic profile that we believe is consistent with previous findings suggesting that the large metabolomic difference is primarily driven by the post eclosion status of these samples, which is likely to override the aging signal of interest, as this status is unique to these samples and has no representative in the C-type samples (Erkosar et al. 2024).

###### Within-regime prediction

For the within-regime predictions, we used an iterative leave-one-replicate-out (LORO) strategy on each regime separately. At each iteration of the LORO procedure, all samples from one replicate were withheld. This left 36 C-type samples for training the within-regime C-type clocks and n=28 A-type samples for the within-regime A-type clocks. This was done to avoid analysis of age predictions on biological replicates that were also present in the training data. See Figure S4 for an example of this phenomenon where the training data has an *R^2^* predictive accuracy of 0.997 while the test data has an *R^2^* predictive accuracy of 0.832. At each iteration, 5-fold CV elastic net models were trained in the glmnet package in R (Friedman et al 2023), to predict age using the metabolome data of the remaining nine replicates. The predictions from training were not used in the final analysis, only to tune the model during 5-fold CV. During model training, the elastic net penalty parameters L1 and L2 were selected from the default tuning grid in glmnet as those that gave the lowest root mean squared error (RMSE). The model from each LORO iteration was then used to predict the ages of the held-out replicate, and these predictions were used in the clock analysis presented in the main text. This procedure was repeated until all replicates had age predictions from a model that was trained on the data from the remaining replicates within the same regime.

###### Between-regime prediction

For the between-regime predictions, 5-fold CV elastic net models were trained on all data (n=40 C-type samples and n=36 A-type samples) and ages within each regime. The between-regime predictions that were analyzed in the main text were then made by the between-regime clock predicting the ages of the opposite regime. So, together, the within-regime and between-regime strategies resulted in predictions made on data that were not a part of model training.

To compare the feature importance in the between-regime A-type and C-type clocks, we used the coefficients (β) fit to each metabolite in the respective clock models. For both clocks, the elastic net penalty L1 at the lowest RMSE was >0, and so some metabolites have β=0 and are not features in the respective clocks.

We tested the accuracy of the within-regime predictions with the coefficient of determination, *R^2^* (Kvalseth 1985; Alexander et al. 2015). Where *y_i_*(*i*= 1, …, *n*) are the ages of the test set, and 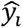 are the predicted ages. 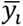 is the mean of *y_i_*, then *R^2^*is calculated as:

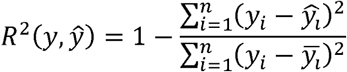

Where an R^2^ closer to 1 represents a better fitting model.

To test our hypothesis that between-regime predictions will reflect differences in the apparent rate of aging within the metabolome, we used a linear model to compare the relationship between ages predicted from the within-regime model over actual age, to the predictions from the between-regime model over age:

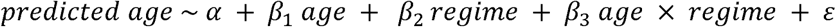

Where α is the intercept of the within-regime model, β*_1_* is the slope of within-regime predictions over age, β*_2_* is the effect of between-regime prediction on the intercept, and β*_3_* is the difference in slopes of the between-regime and the within-regime models.

#### Objective 3: Metabolomic Trajectories of Rapid Aging

##### Metabolomic Differentiation Between Selection Regimes

A LMM was used to compare the metabolite levels between populations under different selection regimes, either A-type or C-type, over time to test for the effects of selection, age, and the interaction between selection and age on the normalized abundance data:

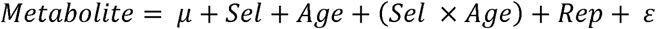

Where selection regime (*Sel*) and age from egg (*Age*) were treated as fixed effects and replicate population (*Rep*) a random effect. The day 9 A-type samples were also withheld from this analysis, as we believe that the large metabolomic difference present for this group of samples is primarily driven by the post eclosion status of these samples, which is likely to override the aging signal of interest and has no equivalent in the C-type samples. We corrected for false discovery using Benjamini Hochberg (1995) approach with a corrected significance threshold of FDR < 0.01. This model was compared to a larger model that accounted for differences between the long-standing populations, and the more recently derived populations (e.g. ACO, AO, etc.), using the Akaike information criterion (AIC). However, adding this term did not improve the fit of the model (see Table S7 for results). The results of the LMM were visualized as an UpSet plot made using the UpSetR package (1.4.0) in R (Conway et al. 2017). The UpSet plot depicts the number of metabolites that were significant for each of the regression terms (FDR < 0.01), as well as the number of significant metabolites shared between any combination of the terms.

##### Characterizing Metabolite Abundance Patterns

To characterize metabolite abundance patterns associated with the accelerated aging phenotype observed from the A-type selection regime, we compared normalized metabolite abundance patterns for different age classes between selection regimes. We calculated mean normalized abundance of each of the 202 metabolites detected at each age within each selection regime (Table S8). We then used the pheatmap package in R (Kolde 2022) to generate a heatmap with hierarchical clustering based on the Euclidean distance and complete linkage to organize both rows (metabolites) and columns (selection regime and age) by similarity in the patterns of metabolite abundance. To determine the optimal number of clusters, we performed K-means clustering for a range of K values (1 to 200), calculating the total within-cluster sum of squares (WSS) for each K using 25 initializations. The difference between consecutive WSS values was calculated to assess the rate of change, and the second derivative of the WSS was computed by calculating the difference between consecutive differences. The optimal K (K = 25) was identified as the value corresponding to the largest second derivative, which signifies the “elbow” of the curve, indicating the point where increasing K no longer substantially reduces the WSS (Figure S5).

The functional enrichment of KEGG network nodes by the metabolites in each cluster that contained at least four metabolites with KEGG IDs was assessed using the network diffusion method in the FELLA package in R (Picart-Armada et al. 2018). We ran 10^5^ permutations to derive an empirical P value (p.score) and pathways that were significant (FDR <0.05) were ranked by p.score within each cluster to identify candidate pathways whose activity could explain the metabolite variation in each cluster (Table S6).

## Data Availability

Raw metabolomic and mortality data are available through Dryad (https://doi.org/10.5061/dryad.1ns1rn92x) and scripts used to process and analyze data are available through Github (https://github.com/mphillips67/A-and-C-Metabolomic-Trajectory-Project).

## Acknowledgments

This work was supported by the following: the Instrumentation Grant of the National Institutes of Health (NIH S10 OD021562), by the Nathan Shock Center for Excellence in Basic Biology of Aging (NIH P30 AG013280), National Institutes of Health Maximizing Investigators’ Research Award to MAP (NIH R35GM155286), the UNCF/Bristol-Myers Squibb E.E. Just Faculty Fund, Career Award at the Scientific Interface (CASI Award) from Burroughs Welcome Fund (BWF) ID # 1021868.01, BWF Ad-hoc Award, NIH Small Research Pilot Subaward to 5R25HL106365-12 from the National Institutes of Health PRIDE Program, DK020593, Vanderbilt Diabetes and Research Training Center for DRTC Alzheimer’s Disease Pilot & Feasibility Program, CZI Science Diversity Leadership grant number 2022-253529 from the Chan Zuckerberg Initiative DAF, an advised fund of Silicon Valley Community Foundation to A.H.J. The Howard Hughes Medical Institute Hanna H. Gray Fellows Program Faculty Phase (Grant# GT15655 awarded to MRM), and the Burroughs Welcome Fund PDEP Transition to Faculty (Grant# 1022604 awarded to MRM). We thank Dr. Michael Rose at UC Irvine for sharing his experimentally evolved fruit fly populations.

## Author Contributions

M.A.P. conceptualized and oversaw the project. M.A.P., K.R.A., and Z.G.G. designed the experiment. A.P., R.D.R, and V.V.C. collected all fly samples for metabolomics and carried out mortality assays under the supervision of K.R.A. D.D. and D.R. processed samples for metabolomic characterizations. T.B.B. analyzed mortality data. D.L.H, Z.G.G., and B.R.H. analyzed the metabolomic data. D.L.H., A.H. Jr., Z.V., and M.R.M developed possible functional interpretation of metabolomic results. D.L.H., M.A.P, and B.R.H. wrote the manuscript.

## References

Alexander, D. L. J., Tropsha, A., & Winkler, D. A. (2015). Beware of R2: Simple, Unambiguous Assessment of the Prediction Accuracy of QSAR and QSPR Models. Journal of Chemical Information and Modeling, 55(7). 10.1021/acs.jcim.5b00206

Avanesov, A. S., Ma, S., Pierce, K. A., Yim, S. H., Lee, B. C., Clish, C. B., & Gladyshev, V. N. (2014). Age- and diet-associated metabolome remodeling characterizes the aging process driven by damage accumulation. ELife, 3. 10.7554/elife.02077

Barghi, N., Tobler, R., Nolte, V., Jakšić, A. M., Mallard, F., Otte, K. A., Dolezal, M., Taus, T., Kofler, R., & Schlötterer, C. (2019). Genetic redundancy fuels polygenic adaptation in Drosophila. PLoS Biology, 17(2). 10.1371/journal.pbio.3000128

Barter, T. T., Greenspan, Z. S., Phillips, M. A., Mueller, L. D., Rose, M. R., & Ranz, J. M. (2019). Drosophila transcriptomics with and without ageing. Biogerontology, 20(5). 10.1007/s10522-019-09823-4

Bates, D., Mächler, M., Bolker, B. M., & Walker, S. C. (2015). Fitting linear mixed-effects models using lme4. Journal of Statistical Software, 67(1). 10.18637/jss.v067.i01

Benjamini, Y., & Hochberg, Y. (1995). Controlling the False Discovery Rate: A Practical and Powerful Approach to Multiple Testing. Journal of the Royal Statistical Society: Series B (Methodological), 57(1). 10.1111/j.2517-6161.1995.tb02031.x

Bujak, R., Daghir-Wojtkowiak, E., Kaliszan, R., & Markuszewski, M. J. (2016). PLS-based and regularization-based methods for the selection of relevant variables in non-targeted metabolomics data. Frontiers in Molecular Biosciences, 3(JUL). 10.3389/fmolb.2016.00035

Burke, M. K., Barter, T. T., Cabral, L. G., Kezos, J. N., Phillips, M. A., Rutledge, G. A., Phung, K. H., Chen, R. H., Nguyen, H. D., Mueller, L. D., & Rose, M. R. (2016). Rapid divergence and convergence of life-history in experimentally evolved Drosophila melanogaster. Evolution; International Journal of Organic Evolution, 70(9). 10.1111/evo.13006

Burke, M. K., Dunham, J. P., Shahrestani, P., Thornton, K. R., Rose, M. R., & Long, A. D. (2010). Genome-wide analysis of a long-term evolution experiment with Drosophila. Nature, 467(7315). 10.1038/nature09352

Carnes, M. U., Campbell, T., Huang, W., Butler, D. G., Carbone, M. A., Duncan, L. H., Harbajan, S. v., King, E. M., Peterson, K. R., Weitzel, A., Zhou, S., & Mackay, T. F. C. (2015). The genomic basis of postponed senescence in Drosophila melanogaster. PLoS ONE, 10(9). 10.1371/journal.pone.0138569

Cavigliasso, F., Savary, L., Spangenberg, J. E., Gallart-Ayala, H., Ivanisevic, J., & Kawecki, T. J. (2023). Experimental evolution of metabolism under nutrient restriction: enhanced amino acid catabolism and a key role of branched-chain amino acids. Evolution Letters, 7(4). 10.1093/evlett/qrad018

Charlesworth, B. (1980). Evolution in age-structured populations. Evolution in Age-Structured Populations. 10.2307/4092

Chippindale, A. K., Alipaz, J. A., Chen, H. W., & Rose, M. R. (1997). Experimental evolution of accelerated development in Drosophila. 1. Developmental speed and larval survival. Evolution, 51(5). 10.1111/j.1558-5646.1997.tb01477.x

Conway, J. R., Lex, A., & Gehlenborg, N. (2017). UpSetR: An R package for the visualization of intersecting sets and their properties. Bioinformatics, 33(18). 10.1093/bioinformatics/btx364

Dettmer, K., Aronov, P. A., & Hammock, B. D. (2007). Mass spectrometry-based metabolomics. In Mass Spectrometry Reviews (Vol. 26, Issue 1). 10.1002/mas.20108

Erkosar, B., Dupuis, C., Savary, L., & Kawecki, T. J. (2024). Shared genetic architecture links energy metabolism, behavior and starvation resistance along a power-endurance axis. *Evolution Letters*, qrae062.

Fabian, D. K., Garschall, K., Klepsatel, P., Santos-Matos, G., Sucena, É., Kapun, M., Lemaitre, B., Schlötterer, C., Arking, R., & Flatt, T. (2018). Evolution of longevity improves immunity in Drosophila. In Evolution Letters (Vol. 2, Issue 6). 10.1002/evl3.89

Friedman, J., Hastie, T., Tibshirani, R., Balasubramanian Narasimhan, Kenneth Tay, Noah Simon, Junyang Qian, & James Yang. (2023). Package ‘glmnet’ - Lasso and Elastic-Net Regularized Generalized Linear Models. In CRAN.

Graves, J. L., Hertweck, K. L., Phillips, M. A., Han, M. v., Cabral, L. G., Barter, T. T., Greer, L. F., Burke, M. K., Mueller, L. D., Rose, M. R., & Singh, N. (2017). Genomics of parallel experimental evolution in drosophila. Molecular Biology and Evolution, 34(4). 10.1093/molbev/msw282

Greenspan, Z., Barter, T., Phillips, M., Ranz, J., Rose, M., & Mueller, L. (2024). Genomewide architecture of adaptation in experimentally evolved Drosophila characterized by widespread pleiotropy. Journal of Genetics, 103(1), 8.

Hardy, C. M., Burke, M. K., Everett, L. J., Han, M. v., Lantz, K. M., & Gibbs, A. G. (2018). Genome-Wide Analysis of Starvation-Selected Drosophila melanogaster-A Genetic Model of Obesity. Molecular Biology and Evolution, 35(1). 10.1093/molbev/msx254

Harrison, B. R., Wang, L., Gajda, E., Hoffman, E. v., Chung, B. Y., Pletcher, S. D., Raftery, D., & Promislow, D. E. L. (2020). The metabolome as a link in the genotype-phenotype map for peroxide resistance in the fruit fly, Drosophila melanogaster. BMC Genomics, 21(1). 10.1186/s12864-020-6739-1

Harrison, B. R., Hoffman, J. M., Samuelson, A., Raftery, D., & Promislow, D. E. L. (2022). Modular Evolution of the Drosophila Metabolome. Molecular Biology and Evolution, 39(1). 10.1093/molbev/msab307

Heinrichsen, E. T., Zhang, H., Robinson, E., Ngo, J., Diop, S., Bodmer, R., Joiner, J., Metallo, M., & Haddad, G. G. (2014). Metabolic and transcriptional response to a high-fat diet in Drosophila melanogaster. Molecular Metabolism, 3(1). 10.1016/j.molmet.2013.10.003

Hoffman, J. M., Soltow, Q. A., Li, S., Sidik, A., Jones, D. P., & Promislow, D. E. L. (2014). Effects of age, sex, and genotype on high-sensitivity metabolomic profiles in the fruit fly, Drosophila melanogaster. Aging Cell, 13(4). 10.1111/acel.12215

[dataset]* Hubert DL, et al. 2024. Data from: Selection for early reproduction leads to accelerated aging and extensive metabolic remodeling in Drosophila melanogaster. Dryad. Dataset. 10.5061/dryad.1ns1rn92x

Ives, P. T. (1970). Further Genetic Studies of the South Amherst Population of Drosophila melanogaster. Evolution, 24(3). 10.2307/2406830

Jin, K., Wilson, K. A., Beck, J. N., Nelson, C. S., Brownridge, G. W. G. W., Harrison, B. R., Djukovic, D., Raftery, D., Brem, R. B., Yu, S., Drton, M., Shojaie, A., Kapahi, P., & Promislow, D. (2020). Genetic and metabolomic architecture of variation in diet restriction-mediated lifespan extension in Drosophila. PLoS Genetics, 16(7). 10.1371/journal.pgen.1008835

Jylhävä, J., Pedersen, N. L., & Hägg, S. (2017). Biological Age Predictors. In EBioMedicine (Vol. 21). 10.1016/j.ebiom.2017.03.046

Kezos, J. N., Barter, T. T., Phillips, M. A., Cabral, L. G., Greenspan, Z. S., Arnold, K. R., Azatian, G., Buenrostro, J., Bhangoo, P. S., Khong, A., Reyes, G. T., Rahman, A., Humphrey, L. A., Bradley, T. J., Mueller, L. D., & Rose, M. R. (2023). Building Bridges from Genome to Physiology Using Machine Learning and Drosophila Experimental Evolution. Physiological and Biochemical Zoology, 96(3). 10.1086/724827

Kezos, J. N., Phillips, M. A., Thomas, M. D., Ewunkem, A. J., Rutledge, G. A., Barter, T. T., Santos, M. A., Wong, B. D., Arnold, K. R., Humphrey, L. A., Yan, A., Nouzille, C., Sanchez, I., Cabral, L. G., Bradley, T. J., Mueller, L. D., Graves, J. L., & Rose, M. R. (2019). Genomics of Early Cardiac Dysfunction and Mortality in Obese Drosophila melanogaster. Physiological and Biochemical ZoologyL: PBZ, 92(6). 10.1086/706099

Kolde, R. (2022). Package “pheatmap”: Pretty heatmaps. R Package.

Koliada, A., Gavrilyuk, K., Burdylyuk, N., Strilbytska, O., Storey, K. B., Kuharskii, V., Lushchak, O., & Vaiserman, A. (2020). Mating status affects Drosophila lifespan, metabolism and antioxidant system. Comparative Biochemistry and Physiology -Part AL: Molecular and Integrative Physiology, 246. 10.1016/j.cbpa.2020.110716

Kosakamoto, H., Okamoto, N., Aikawa, H., Sugiura, Y., Suematsu, M., Niwa, R., Miura, M., & Obata, F. (2022). Sensing of the non-essential amino acid tyrosine governs the response to protein restriction in Drosophila. Nature Metabolism, 4(7). 10.1038/s42255-022-00608-7

Kurup, K., Matyi, S., Giles, C. B., Wren, J. D., Jones, K., Ericsson, A., Raftery, D., Wang, L., Promislow, D., Richardson, A., & Unnikrishnan, A. (2021). Calorie restriction prevents age-related changes in the intestinal microbiota. Aging, 13(5). 10.18632/aging.202753

Kvalseth, T. O. (1985). Cautionary note about r2. American Statistician, 39(4). 10.1080/00031305.1985.10479448

Lassen, J. K., Wang, T., Nielsen, K. L., Hasselstrøm, J. B., Johannsen, M., & Villesen, P. (2023). Large-Scale metabolomics: Predicting biological age using 10,133 routine untargeted LC–MS measurements. Aging Cell, 22(5). 10.1111/acel.13813

Laye, M. J., Tran, V., Jones, D. P., Kapahi, P., & Promislow, D. E. L. (2015). The effects of age and dietary restriction on the tissue-specific metabolome of Drosophila. Aging Cell, 14(5). 10.1111/acel.12358

Lu, A. T., Fei, Z., Haghani, A., Robeck, T. R., Zoller, J. A., Li, C. Z., Lowe, R., Yan, Q., Zhang, J., Vu, H., Ablaeva, J., Acosta-Rodriguez, V. A., Adams, D. M., Almunia, J., Aloysius, A., Ardehali, R., Arneson, A., Baker, C. S., Banks, G., … Horvath, S. (2023). Universal DNA methylation age across mammalian tissues. Nature Aging, 3(9). 10.1038/s43587-023-00462-6

Luckinbill, L. S., Arking, R., Clare, M. J., Cirocco, W. C., & Buck, S. A. (1984). Selection for Delayed Senescence in Drosophila melanogaster. Evolution, 38(5). 10.2307/2408433

Meador, J. P., Bettcher, L. F., Ellenberger, M. C., & Senn, T. D. (2020). Metabolomic profiling for juvenile Chinook salmon exposed to contaminants of emerging concern. Science of the Total Environment, 747. 10.1016/j.scitotenv.2020.141097

Meyer, D. H., & Schumacher, B. (2024). Aging clocks based on accumulating stochastic variation. Nature Aging. 10.1038/s43587-024-00619-x

Parrott, B. B., & Bertucci, E. M. (2019). Epigenetic Aging Clocks in Ecology and Evolution. In Trends in Ecology and Evolution (Vol. 34, Issue 9). 10.1016/j.tree.2019.06.008

Patti, G. J., Yanes, O., & Siuzdak, G. (2012). Innovation: Metabolomics: the apogee of the omics trilogy. In Nature Reviews Molecular Cell Biology (Vol. 13, Issue 4). 10.1038/nrm3314

Phillips, M. A., Arnold, K. R., Vue, Z., Beasley, H. K., Garza-Lopez, E., Marshall, A. G., Morton, D. J., McReynolds, M. R., Barter, T. T., & Hinton, A. (2022). Combining Metabolomics and Experimental Evolution Reveals Key Mechanisms Underlying Longevity Differences in Laboratory Evolved Drosophila melanogaster Populations. International Journal of Molecular Sciences, 23(3). 10.3390/ijms23031067

Phillips, M. A., Long, A. D., Greenspan, Z. S., Greer, L. F., Burke, M. K., Villeponteau, B., Matsagas, K. C., Rizza, C. L., Mueller, L. D., & Rose, M. R. (2016). Genome-wide analysis of long-term evolutionary domestication in Drosophila melanogaster. Scientific Reports, 6. 10.1038/srep39281

Picart-Armada, S., Fernández-Albert, F., Vinaixa, M., Yanes, O., & Perera-Lluna, A. (2018). FELLA: An R package to enrich metabolomics data. BMC Bioinformatics, 19(1). 10.1186/s12859-018-2487-5

Rodrigues, M. A., Dauphin-Villemant, C., Paris, M., Kapun, M., Mitchell, E. D., Kerdaffrec, E., & Flatt, T. (2023). Germline proliferation trades off with lipid metabolism in *Drosophila*. Evolution Letters. 10.1093/evlett/qrad059

Rose, M., & Charlesworth, B. (1980). A test of evolutionary theories of senescence. Nature, 287(5778). 10.1038/287141a0

Rose, M., Passananti, H., & Matos, M. (2010). Methuselah Flies - A Case Study in the Evolution of Aging. In Methuselah Flies - A Case Study in the Evolution of Aging. 10.1142/9789812567222

Rose, M. R. (1984). Laboratory Evolution of Postponed Senescence in Drosophila melanogaster. Evolution, 38(5). 10.2307/2408434

Rose, M. R., & Charlesworth, B. (1981). Genetics of life history in Drosophila melanogaster. II. Exploratory selection experiments. Genetics, 97(1). 10.1093/genetics/97.1.187

Rose, M. R., Rauser, C. L., Benford, G., Matos, M., & Mueller, L. D. (2007). Hamilton’s forces of natural selection after forty years. In Evolution (Vol. 61, Issue 6). 10.1111/j.1558-5646.2007.00120.x

Rutledge, J., Oh, H., & Wyss-Coray, T. (2022). Measuring biological age using omics data. In Nature Reviews Genetics (Vol. 23, Issue 12). 10.1038/s41576-022-00511-7

Team, R. C. (2023). R Core Team 2023 R: A language and environment for statistical computing. R foundation for statistical computing. https://www.R-project.org/. R Foundation for Statistical Computing.

Tobler, R., Hermisson, J., & Schlötterer, C. (2015). Parallel trait adaptation across opposing thermal environments in experimental Drosophila melanogaster populations. Evolution, 69(7). 10.1111/evo.12705

Turner, T. L., Stewart, A. D., Fields, A. T., Rice, W. R., & Tarone, A. M. (2011). Population-based resequencing of experimentally evolved populations reveals the genetic basis of body size variation in Drosophila melanogaster. PLoS Genetics, 7(3). 10.1371/journal.pgen.1001336

Wang, R., Yin, Y., Li, J., Wang, H., Lv, W., Gao, Y., Wang, T., Zhong, Y., Zhou, Z., Cai, Y., Su, X., Liu, N., & Zhu, Z. J. (2022). Global stable-isotope tracing metabolomics reveals system-wide metabolic alternations in aging Drosophila. Nature Communications, 13(1). 10.1038/s41467-022-31268-6

Wickham, H. (2016). ggplot2 Elegant Graphics for Data Analysis (Use R!). Springer.

Zhao, X., Golic, F. T., Harrison, B. R., Manoj, M., Hoffman, E. v., Simon, N., Johnson, R., MacCoss, M. J., McIntyre, L. M., & Promislow, D. E. L. (2022). The metabolome as a biomarker of aging in Drosophila melanogaster. Aging Cell, 21(2). 10.1111/acel.13548

Zou, H., & Hastie, T. (2005). Regularization and variable selection via the elastic net. Journal of the Royal Statistical Society. Series B: Statistical Methodology, 67(2). 10.1111/j.1467-9868.2005.00503.x

