## Supplemental Figures for "Selection for early reproduction leads to accelerated aging and extensive metabolic remodeling in *Drosophila melanogaster*"

**Supplementary Figures**

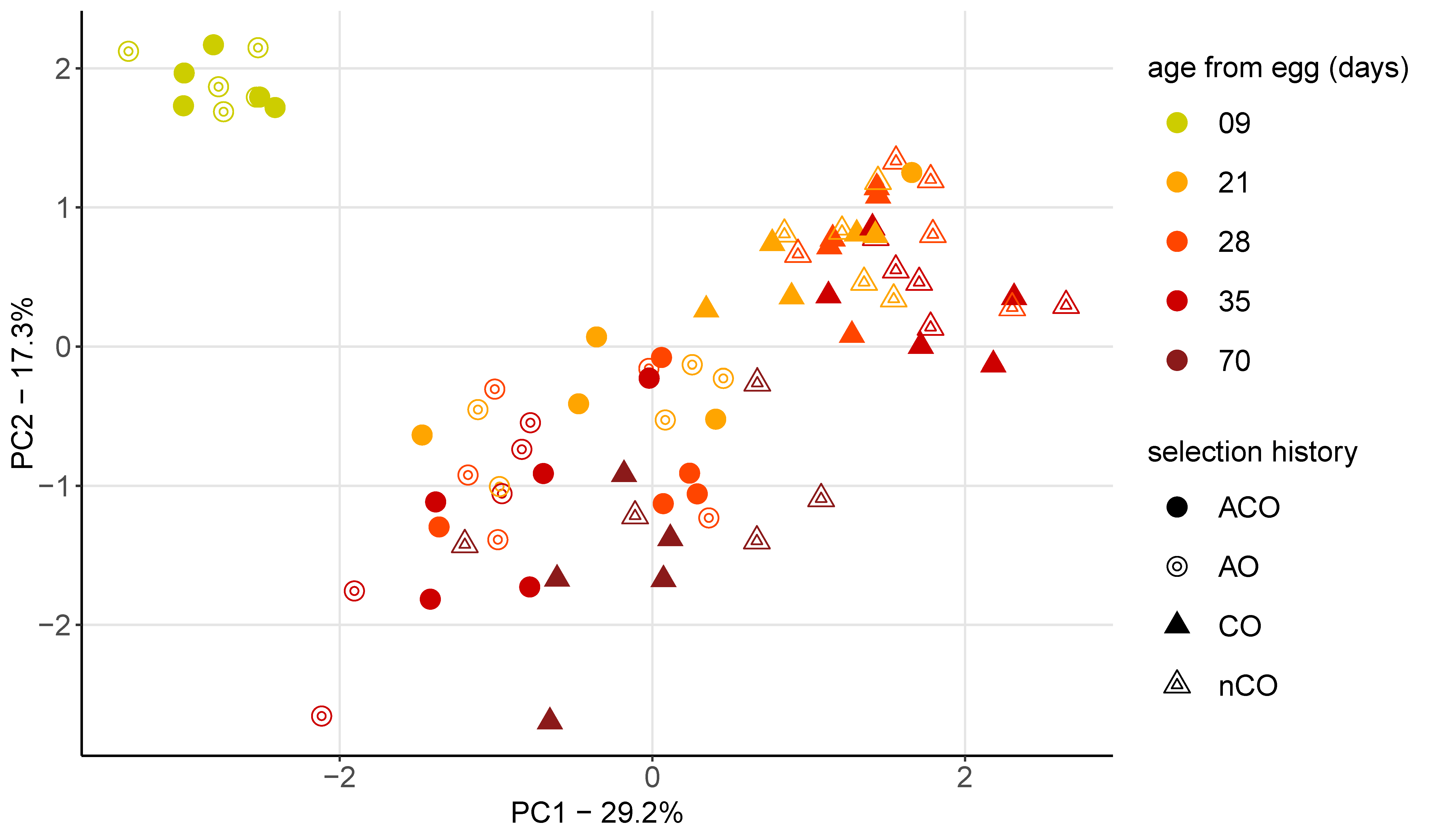

**Supplementary Figure S1.** Principal component analysis (PCA) plot shows how mean centered metabolite abundance clusters by age from egg in days (color), selection regime (shape), and selection history (shape density) along the first principal component (PC1) and the second (PC2). Populations from the A-type selection regime (circles), and C-type selection regime (triangles) are differentiated as having either a long-standing selection history (solid fill) or a recently derived selection history (hollow).

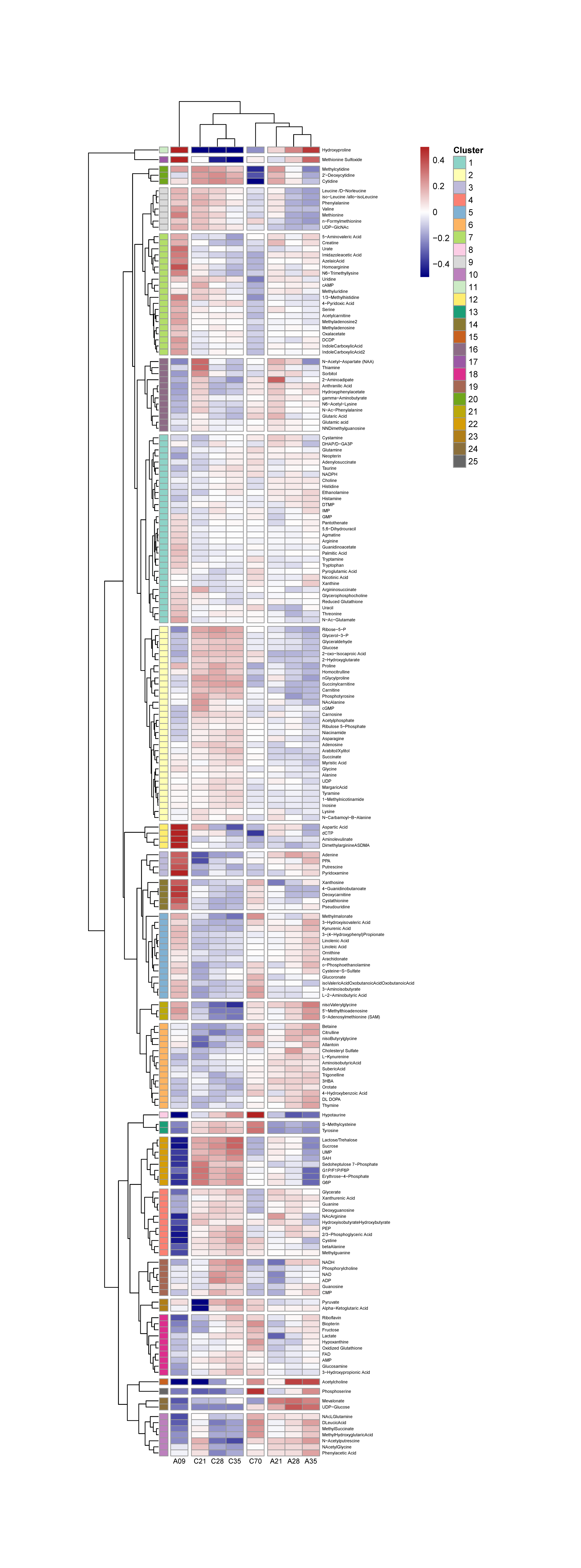

**Supplementary Figure S2:** **Hierarchal clustering of mean normalized metabolite abundance clustered by regime and age.** Cells represent mean normalized metabolite abundance for each of 10 replicates for each selection regime and age (in days since egg) for 176 metabolites that were significant for any term from LMM (Table S5). Hierarchical clustering based on the Euclidean distance and complete linkage to organize both rows (metabolites) and columns (selection regime and age) by similarity in the patterns of metabolite abundance. Red color indicates a higher mean normalized abundance, and blue color indicates a lower mean normalized abundance.

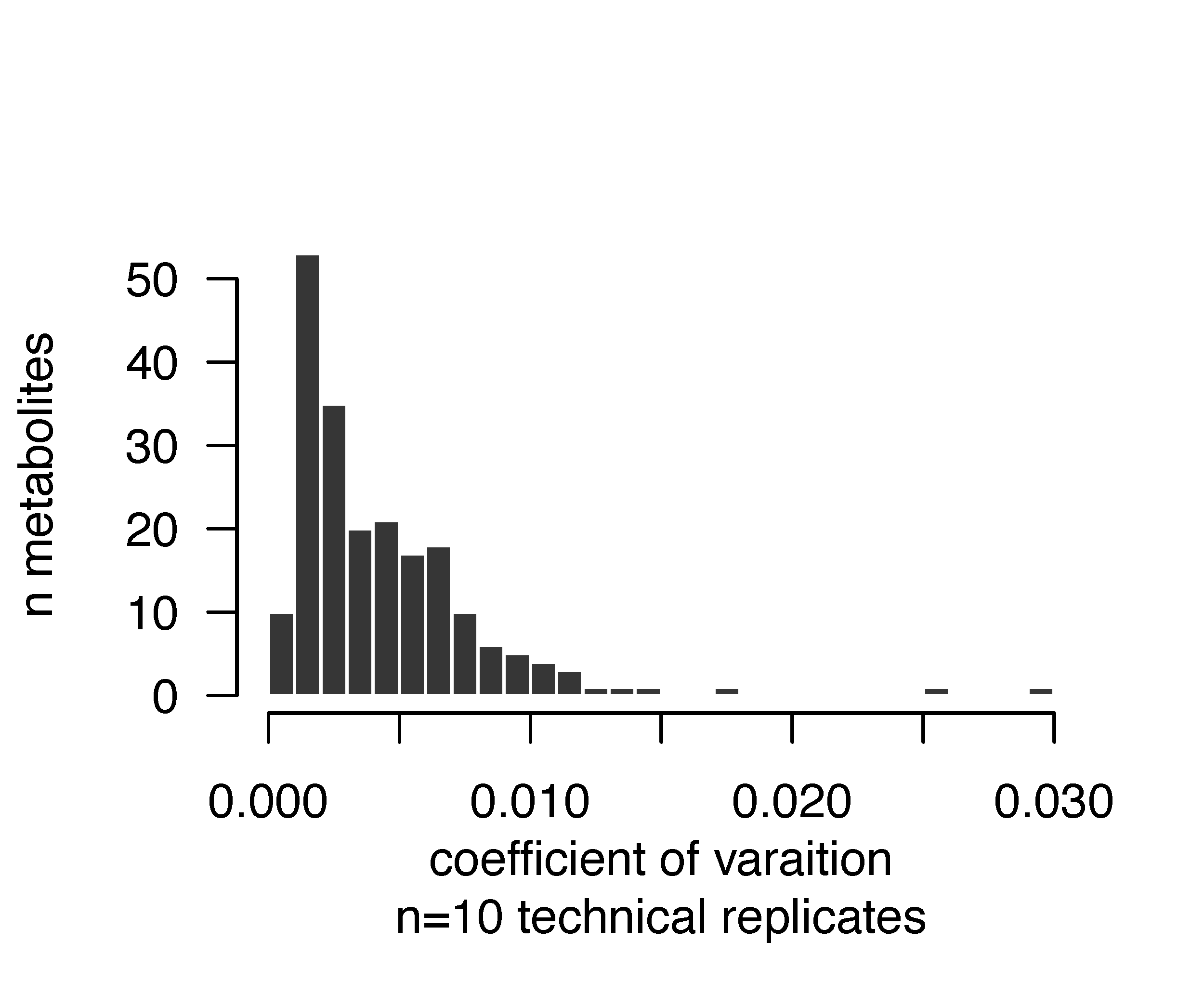

**Supplementary Figure S3. Reproducibility among the 202 metabolites on the LC-MS.** A histogram of the coefficient of variation (CV = SD of log-metabolite values / mean log-metabolite value) for the 202 targeted metabolites detected in all samples. Mean CV=0.0044, with a range of 6x10^-4^ to 0.029. The CVs reported here are from a single, pooled liquid sample of all 80 biological samples run in the analysis. This pooled sample was resampled 10 times and run interspersed among the study samples in the LC-MS. These 10 technical samples reflect the reproducibility of detection withing each of the 202 metabolites across the entire analysis.

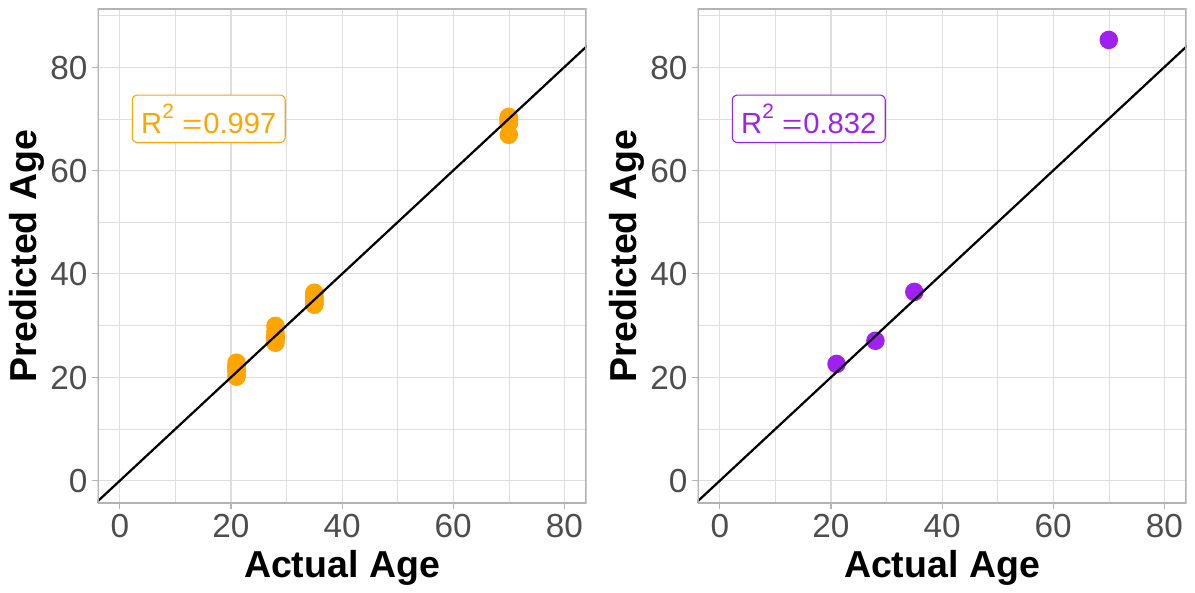

**Supplementary Figure S4. One Singular Elastic Net Regression Run:** Predicted age from egg (days) vs chronological age from egg (days) for nine C-type populations (orange) and the CO3 (purple) population based on elastic net models. The left panel shows the predictions from metabolomic profiles for the populations the model was trained on while the right panel shows the predictions from the metabolomic profiles for the left-over population the model was tested on. We plot the left-out population in our main text to avoid overfitting.

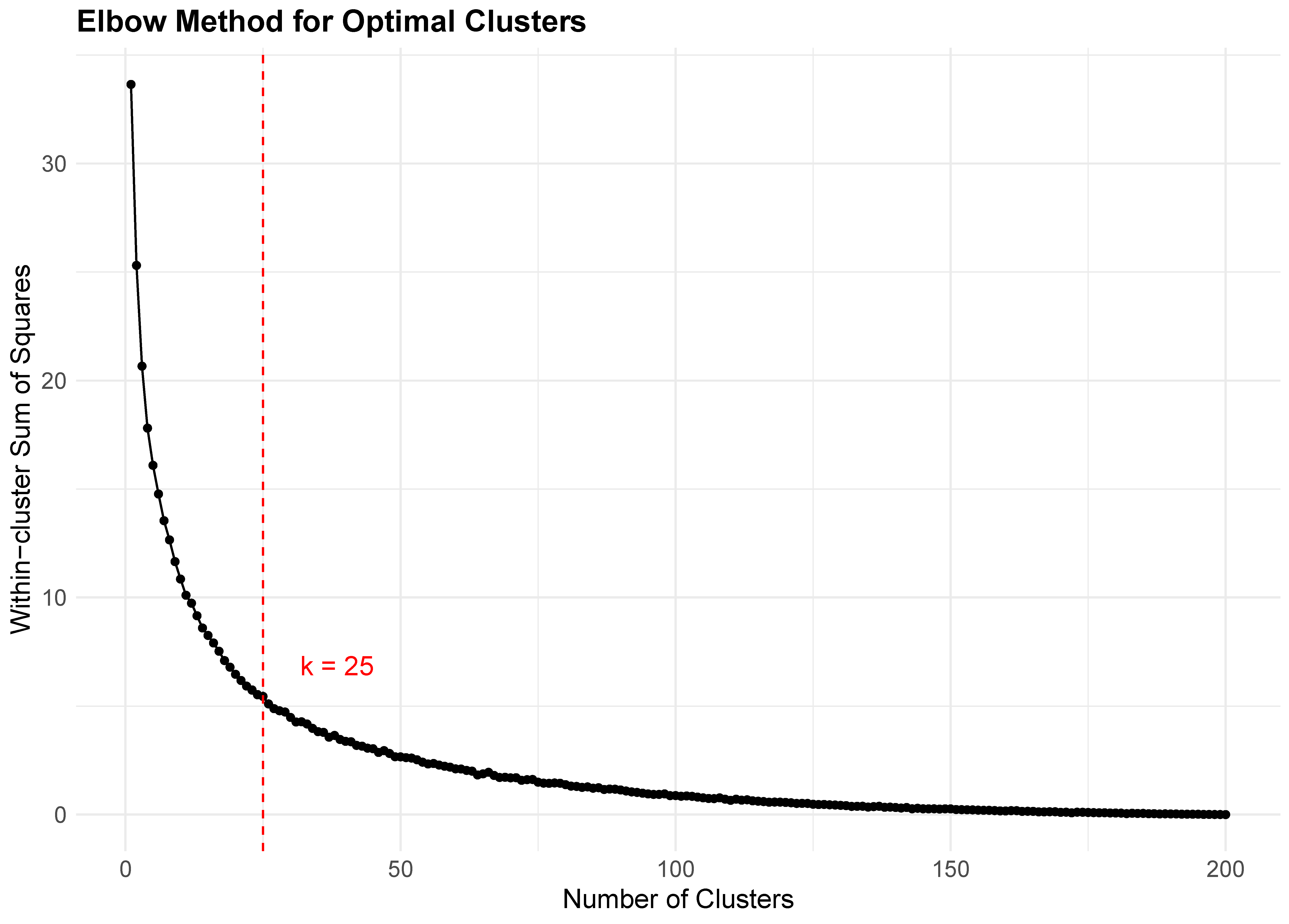

**Figure S5. Elbow method for determining the optimal number of clusters (K)** used for hierarchical clustering of mean normalized metabolite abundance (Figures 8 and S2). The within-cluster sum of squares (WSS) was calculated for K values ranging from 1 to 200. The first and second differences of WSS were computed to identify the point where the rate of decrease slows the most, corresponding to the 'elbow' in the curve. The optimal K (25) was determined as the value where the second derivative of WSS is minimized, indicating the best balance between model complexity and clustering efficiency.
